# PERK-dependent reciprocal crosstalk between ER and non-centrosomal microtubules coordinates ER architecture and cell shape

**DOI:** 10.1101/2021.01.19.426991

**Authors:** Miguel Sánchez-Álvarez, Fidel Lolo, Heba Sailem, Patricia Pascual-Vargas, Giulio Fulgoni, Mar Arias-García, Miguel Ángel del Pozo, Chris Bakal

## Abstract

The architecture of the endoplasmic reticulum (ER) is tightly controlled as a key determinant of its function. Its dynamics are linked to those of the cytoskeleton, but our understanding of how this coordination occurs and what its functional relevance is, is limited. We found the Unfolded Protein Response (UPR) transducer EIF2AK3/PERK is essential for acute stress-induced peripheral redistribution and remodeling of the ER, through eIF2a phosphorylation and translation initiation shutdown. PERK-mediated eIF2a phosphorylation can be bypassed by blocking ribosome activity; by depleting microtubule-anchoring ER proteins such as REEP4, p180/RRBP1 and Climp63/CKAP4; or by disrupting the microtubule cytoskeleton. Notably, specific disruption of non-centrosomal microtubules, but not centrosome depletion, relieved blockade of ER redistribution in PERK-deficient cells. Conversely, PERK deficiency stabilized non-centrosomal microtubules, promoting polarized protrusiveness in epithelial cells and neuroblasts. We propose that PERK coordinates ER architecture and homeostasis with cell morphogenesis by coupling ER remodeling and non-centrosomal MT dynamics.

## INTRODUCTION

The eukaryotic endoplasmic reticulum (ER) is an intricate system of intracellular membrane domains delimiting a single luminal space, continuous with the outer nuclear envelope. The architecture of the ER and its dynamics contribute to the several essential functions of this organelle, including calcium and redox homeostasis, complex lipid metabolism, management of other endomembrane systems, and the maturation and assisted folding of ~30% of the proteome^1^. ER membrane subdomains can adopt discrete shapes (including ER ‘tubules’ (peripheral, reticular tubes of ER, with rather low densities of associated ribosomes) and ER ‘sheets’ (flat enlargements or “cisterns” of peripheral ER, usually rich in bound polysomes)^2^. The local concentration and activity of “ER shapers” such as reticulons, atlastins and other auxiliary resident proteins^3^ determines the shape of the ER. Accordingly, physical expansion of the ER is an integral component of its adaptive response to functional imbalances (generically termed “ER stress”)^4^. ER expansion is integrated with the Unfolded Protein Response of the ER (UPR^ER^), a surveillance mechanism that continuously gauges ER luminal environment and membrane integrity and engages adaptive programs as required^5–7^.

The UPR^ER^ of higher eukaryotes comprises three main branches, each of them driven by a specific, ER-resident transducer. Inositol Requiring enzyme 1 (IRE1; also known as Endoplasmic Reticulum To Nucleus Signaling 1, ERN1) orchestrates a complex adaptive transcriptional response through the unconventional splicing of *XBP1* (X-box binding protein 1) mRNA, eliciting its translation as a potent transcriptional transactivator of ER chaperones and lipid anabolism^7,8^. Upon activated intramembrane cleavage, Activation Transcription Factor 6 (ATF6) also promotes adaptive transcriptional programs, driving red/ox regulation, ER chaperone expression and lipid metabolism enzymes^7,9^. Both UPR branches are involved in ER membrane *de novo* synthesis and ER physical expansion^10–12^. The third branch is operated by the EIF2A kinase 3/PKR-like ER Kinase (EIF2AK3/PERK), one of the four known stress-associated kinases in metazoans capable of phosphorylating the essential translation regulator eukaryotic initiation factor 2a (eIF2α). eIF2α is a GTPase subunit of the translation initiation ternary complex capped mRNA-ribosomal subunit 60S-eIF^7^; phosphorylation on its conserved Ser51 residue locks eIF2α in its GDP-bound, inactive state and leads to sequestration of the ternary complex from further assembling a processive ribosome. Therefore, eIF2α phosphorylation is a node onto which several stress responses (including UPR through PERK) converge to regulate protein synthesis. Limiting client protein overload in the stressed ER through translation downregulation is thus commonly proposed as the main role of PERK^7,13^. However, PERK-dependent protein translation attenuation is integrated with other functional outputs beyond control of client protein burden on the ER, such as regulation of calcium trafficking and apoptosis^14–16^.

The ER communicates with other membrane-bound organelles, plasma membrane subdomains, and the cytoskeleton^17^. While lower eukaryotes and plants mostly rely on actin cytoskeleton dynamics for the shaping and partition of their ER^18,19^, the metazoan ER is tightly coupled to the microtubule (MT) cytoskeleton^20–23^. Several integral proteins of the ER membrane such as Stromal interaction molecule 1 (STIM1), Ribosome-binding protein 1 (RRBP1/p180), cytoskeleton-linking membrane protein 63 (Climp63/CKAP4), Receptor expression-enhancing proteins 1-4 (REEPs1-4) or reticulons can engage in physical contacts with MTs^24–28^. Extension, fusion and reticulation of ER tubules are guided by dynamic microtubule bundles^17^. ER sheets also establish anchoring interactions with microtubules, and they appear to do so preferentially through p180/RRBP1 and Climp63, which in turn also establish interactions with polysome/translocon complexes enriched in these areas of “rough” ER ^29,30^. Importantly, these interactions are a means by which changes in ER morphology can affect MT organization, and viceversa. For example, p180/RRBP1 overexpression can induce hyperstabilization of microtubule bundles and promote the formation of tight, collapsed structures^25^.

We developed automated image-based RNAi screening procedures to explore the genetic regulation of ER expansion and redistribution upon pharmacologically challenging ER homeostasis. We found the UPR effector PERK is essential for ER redistribution to the cell periphery during ER expansion upon induction of ER stress. This activity is dependent on translation initiation shutdown through eIF2α phosphorylation, and can be bypassed by blocking translation initiation and polysome disassembly, but not by shutting down translation elongation. Combinatorial siRNA screening revealed that depletion of proteins linking microtubules with the ER, such as REEP4, p180/RRBP1 and Climp63/CKAP4, specifically rescued the ER collapse associated with disruption of PERK signaling. Importantly, while centrosome depletion did not have an observable impact on ER architecture dynamics, inhibition of CAMSAP2-stabilized non-centrosomal microtubules promoted the peripheral expansion of ER sheet structures, and fully rescued ER redistribution in PERK-depleted cells. Conversely, abrogation of PERK activity led to phenotypes of increased polarity and low number of large protrusions in epithelial cells, for which epistasis with non-centrosomal microtubules was also observed. Thus, during ER stress PERK inhibits translation initiation at the ribosomes, which alters the coupling of ER to non-centrosomal MTs through specific ER-MT linkers such as CKAP4/Climp63 and RRBP1/p180, and both facilitates expansion and modulates MT cytoskeleton arrangement. Our observations highlight the existence of additional key roles for PERK on cell homeostasis beyond the curbing of ER client protein synthesis.

## RESULTS

### PERK is a novel regulator of ER redistribution during acute ER stress in epithelial cells

With the aim of exploring mechanisms determining ER architecture during ER stress, we developed an image analysis pipeline for automated high-content ER morphology analysis in single cells. (fig. 1A; ^31,32^). An array of features (periphery/perinuclear averaged intensity ratio of the ER; image textures, reflecting different aspects of ER architecture; and subcellular distribution) is extracted from single cells, together with other morphological features of the whole cell (fig. 1A and fig. S1A). To assess the validity of our approach to capture significant changes in ER morphogenesis across different conditions, we first compared wild type cells with cells exposed to the N-glycosylation inhibitor tunicamycin, which provokes acute ER stress and a prominent redistribution and expansion of the ER in epithelial cells, together with changes in image texture (fig. 1A). The ratio of ER signal density on the cell periphery as compared with the perinuclear region was informative of adaption to ER stress (fig. 1A, right panels). We further tested this system by interrogating, both in untreated and tunicamycin-treated cells, a small collection of siRNAs, targeting well-established direct regulators of ER morphogenesis and homeostasis (see Table S1). Notably, knockdown of Inositol Requiring Enzyme 1 alpha (IRE1a), a key regulator of ER membrane expansion^4,33^, led to reduced peripheral ER distribution across conditions (fig. 1B and S1B). Thus, we conclude we are able to quantify physiologically relevant changes in ER morphology in single cells during ER stress.

**Figure 1.**
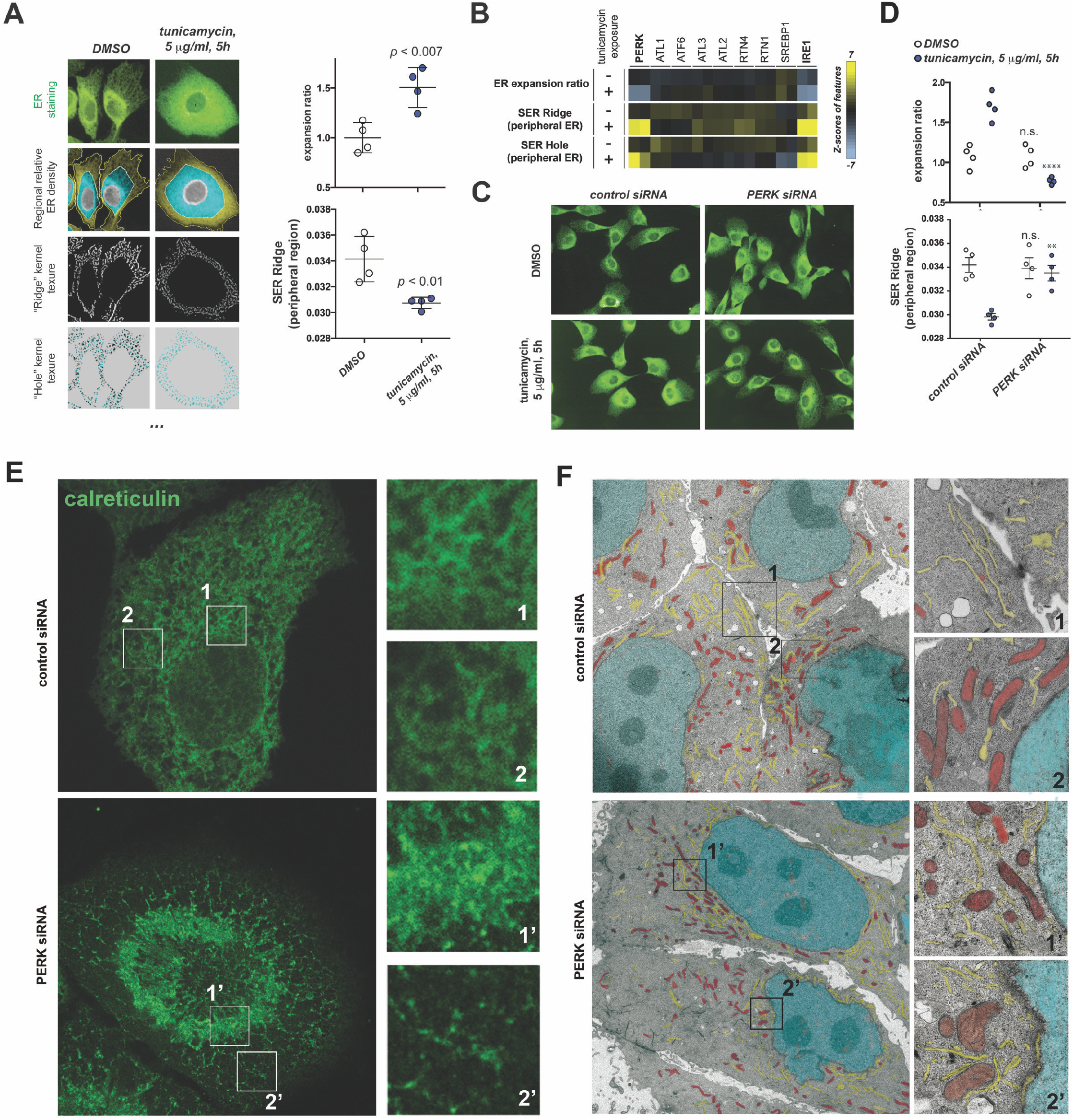
An automated image-based assay identifies EIF2AK3/PERK as required for ER remodeling during acute ER stress. **(A)** Examples of automated image analysis features informative of ER architecture remodeling in MCF10A cells treated as indicated, from unprocessed immunofluorescence images of total ER (anti-calreticulin immunostaining). Graphs (rightmost panels) are derived from four independent replicates of indicated conditions. **(B-D)** Results from automated image-based exploration of genetic regulators of ER stress-driven ER remodeling, using pools of 4 siRNA duplexes to interrogate each chosen gene. [B] Heatmaps (Z-scores) of selected features, informative for ER remodeling, across conditions, as compared to control cells. [C] Representative images for indicated conditions. [D] Graphs derived from well-normalized values for indicated features across conditions (four independent replicates. **(E, F)** STED microscopy (calreticulin immunostaining) [E] and electron microscopy [F] of MCF10A cells exposed for 6h to 5μg/ml of tunicamycin. Magnified cropped images show details of indicated ROIs at perinuclear or peripheral cell regions. Pseudocolouring in [F]: *cyan*-nuclei; *red*-mitochondria; *yellow*-ER. *p* values are indicated across experiments. Statistical significance across assays was assessed by paired t-Student test; *: *p* <0.05; **: *p* <0.01; ***: *p* <0.005. n.s.: *p* >0.05

We found that depletion of the protein kinase R (PKR)-like endoplasmic reticulum kinase (PERK/EIF2AK3), a well-established essential factor for ER homeostasis^13^, led to a marked impairment for ER subcellular redistribution upon exposure to tunicamycin (fig. 1B-D and fig. S1A). Superresolution light microscopy and electron microscopy confirmed that the ER of cells depleted of PERK is not rearranged to increase peripheral sheets in response to ER stress, and collapses at the perinuclear region (fig. 1E and F). To assess that these phenotypes were not due to off-target effects, four different siRNA sequences were chosen for secondary validation. qRT-PCR assays consistently showed a downregulation of at least ~80% in PERK mRNA levels upon transfection of any of the tested siRNA sequences (fig. S1C). Transfection of all four siRNAs recapitulated the phenotype of impaired peripheral ER redistribution upon tunicamycin challenge (fig. 2A, lower row; fig. 2B). PERK knock-down in other cell lines of different origins, such as a transformed MCF10A clone (ATI clone, expressing the constitutively active H-Ras G12V mutant) or two other tumour epithelial cell lines (HeLa and MDA-MB231) resulted in identical phenotypes (fig. S1D and E). Furthermore, the phenotype of impaired ER subcellular redistribution upon acute exposure to tunicamycin could also be followed through live imaging of a stable MCF10A-derived cell line expressing EGFP-tagged Sec61β (fig. S1F).

**Figure 2.**
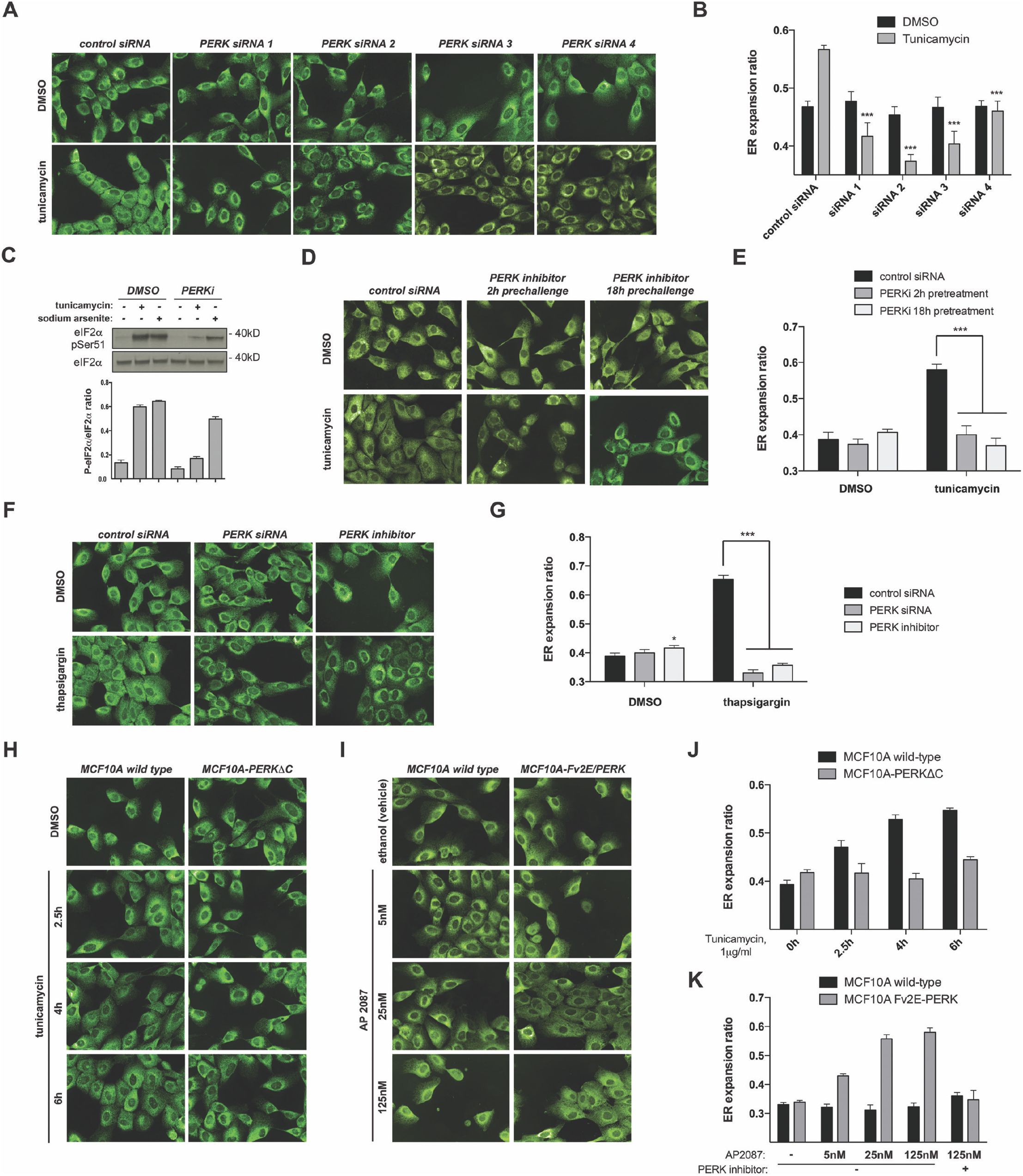
Validation of PERK as an essential effector of acute ER stress-driven ER remodeling. **(A, B)** Independent analysis of ER remodelling phenotypes for each PERK-targeting siRNA duplex. [A] Representative unprocessed, automatically acquired images of anti-calreticulin-immunostained MCF10A cells across indicated conditions. [B] Graphs derived from four independent replicates (~2000 cells per well). **(C-E)** Allosteric inhibition of PERK kinase activity recapitulates the phenotype observed upon siRNA-mediated PERK depletion. [C] Assessment of the activity and specificity of the inhibitory compound (PERKi; GSK2606414, see Matherials and Methods) by western blot analysis of whole-cell lysates across indicated conditions. [D] Representative images (calreticulin immunostaining) and [E] data derived from eight biological replicates (~2000 cells per well) across indicated treatment conditions. **(F-G)** Recapitulation of previous results upon acute ER stress induction using a different mechanism (Ca^2+^ depletion from the ER, using the SERCA2 inhibitor thapsigargin). [F] Representative anti-calreticulin immunofluorescence staining across indicated conditions; [G] data derived from four biological replicates (~2000 cells per well) across indicated treatments. **(H, J)** Stable expression of an inactive PERK-ΔC truncated mutant blunts ER stress-induced PERK signaling and ER peripheral remodelling. Data is derived from eight biological replicates (~2000 cells per well) across indicated treatments. **(I, K)** Exposure of cells expressing a synthetic fusion PERK protein with homodimerizing domains (PERK-Fv2E), to the AP20187 homodimerizer enables for PERK kinase-dependent ER remodelling bypassing ER stress induction. Data is derived from eight biological replicates (~2000 cells per well) across indicated treatments. Statistical significance across assays was assessed by paired t-Student test; *: *p* <0.05; **: *p* <0.01; ***: *p* <0.005. n.s.: *p* >0.05

We further validated our observations using the allosteric PERK kinase inhibitor GSK2606414 (fig. 2C) for different times, previous to tunicamycin challenge. PERK kinase inhibition again recapitulated the ER collapse associated with PERK siRNA depletion (fig. 2D and E). Of note, disruption of PERK kinase activity using this chemical inhibitor in unchallenged cells also led to moderate phenotypic alterations regarding cell elongation and peripheral ER architecture, similar to those observed for PERK siRNA-mediated knock-down. PERK siRNA transfection or exposure to PERK kinase inhibitor were also associated with ER collapse when cells were challenged with a different source of ER stress, such as the sarco/endoplasmic reticulum Ca2+-ATPase inhibitor thapsigargin (fig. 2F and G).

We further tested two established cell models, derived from the MCF10A epithelial line, which reprogram PERK regulation and output. First, we looked into the phenotype of the PERK-ΔC cell line—which expresses a C-terminal truncated mutant with dominant negative properties^34,35^)—across different conditions. The PERK-ΔC-expressing cell line exhibited partially impaired redistribution of their ER upon tunicamycin exposure (fig 2H and J). We studied the behavior of another MCF10A-derived cell line, Fv2E-PERK, which expresses a synthetic PERK construct by which it is possible to uncouple PERK activation from canonical ER stress (^34,35^; fig. S2A). Exposure to the B/B homodimerizer AP20187 provoked apparent ER peripheral expansion in Fv2E-PERK cells, but not in wild-type MCF10A cells (fig. 2I and K). These observations support that the phenotypes we observe are specifically derived from altered PERK kinase activity.

### PERK/EIF2AK3 kinase role in remodeling of peripheral ER architecture depends on eIF2α phosphorylation-driven translation attenuation

In certain cell models, PERK is required for the activation of the lipid anabolism regulator Sterol Regulatory Element Binding Protein 1 (SREBP1; ^36^). Thus we considered deficiency in ER expansion in PERK-deficient cells could be due to impaired lipid anabolism or *de novo* membrane synthesis. However, in our hands, siRNA-mediated depletion of PERK did not impact significantly on the activation of the canonical ER lipid anabolism regulators such as SREBP1, or X-box binding protein 1 mRNA processing (XBP1) (^10–12,36^, fig. S2B). Furthermore, cells depleted for these regulators exhibited ER phenotypes different from those displayed by cells knocked down for PERK (see above Fig. S1B). Importantly, the increase in total ER membrane content upon ER stress induction in PERK-depleted cells is comparable to that of wild-type cells (fig. S2C). Thus, our observations suggested a potential novel role for PERK in the control of ER architecture remodelling and subcellular redistribution during ER stress, distinct from *de novo* ER membrane synthesis.

We next investigated the role of Activated Transcription Factor 4 (ATF4) in the role of PERK-mediated ER expansion. ATF4 is a a key UPR transcriptional regulator whose translation is dependent on the ribosomal frameshift elicited by PERK-dependent eIF2α phosphorylation^15^. PERK depletion in our cell system blunted ER stress-driven ATF4 translation (fig. S2D). But siRNA-mediated depletion of ATF4 did not recapitulate the phenotypic characteristics associated with the abrogation of PERK activity, and in fact led to moderate but significantly opposite effects, regarding ER remodeling, cell spreading and overall morphology across the tested conditions (fig. S2E and F). These observations support the notion that the role of PERK in the phenotypes we observe is not dependent on downstream gene expression programs driven by the ATF4/CHOP axis.

We hypothesized that PERK-dependent translation of targets in addition to ATF4 might directly underlie stress-induced ER redistribution. We thus implemented a multiplexed method to quantify global translation activity on a single-cell basis while simultaneously determining the subcellular distribution and architecture of the ER. Polysomes engaged in active translation are immunolabelled and visualized after a brief pulse of puromycin (puromycylation; hereon termed ‘PMY’ (fig. S3A,^37^). The signal is highly specific because either omission of PMY, or robust disruption of ribosomal assembly through different approaches before exposure to a puromycin pulse, completely abolish immunolabelling (fig. S3B). Acute induction of ER stress leads to a substantial decrease in PMY signal, which correlates with upregulation of phosphor-eIF2α signal, in a PERK-dependent manner (fig. 3A). We plotted ER redistribution measurements against PMY signal across different conditions for single wild-type MCF10A cells (fig. 3B). Importantly, translation activity (as judged by PMY intensity) and degree of relative peripheral ER density exhibited an *inverse* correlation on a single-cell basis. Moreover, phosphor-eIF2α signal (inhibition of translation) followed an opposite (positive) correlation pattern with ER expansion on a single-cell basis (fig. 3B, lower panels). Thus as expected acute induction of ER stress robustly increased ER expansion but suppressed translation globally (fig. 3B, upper panels).

**Figure 3.**
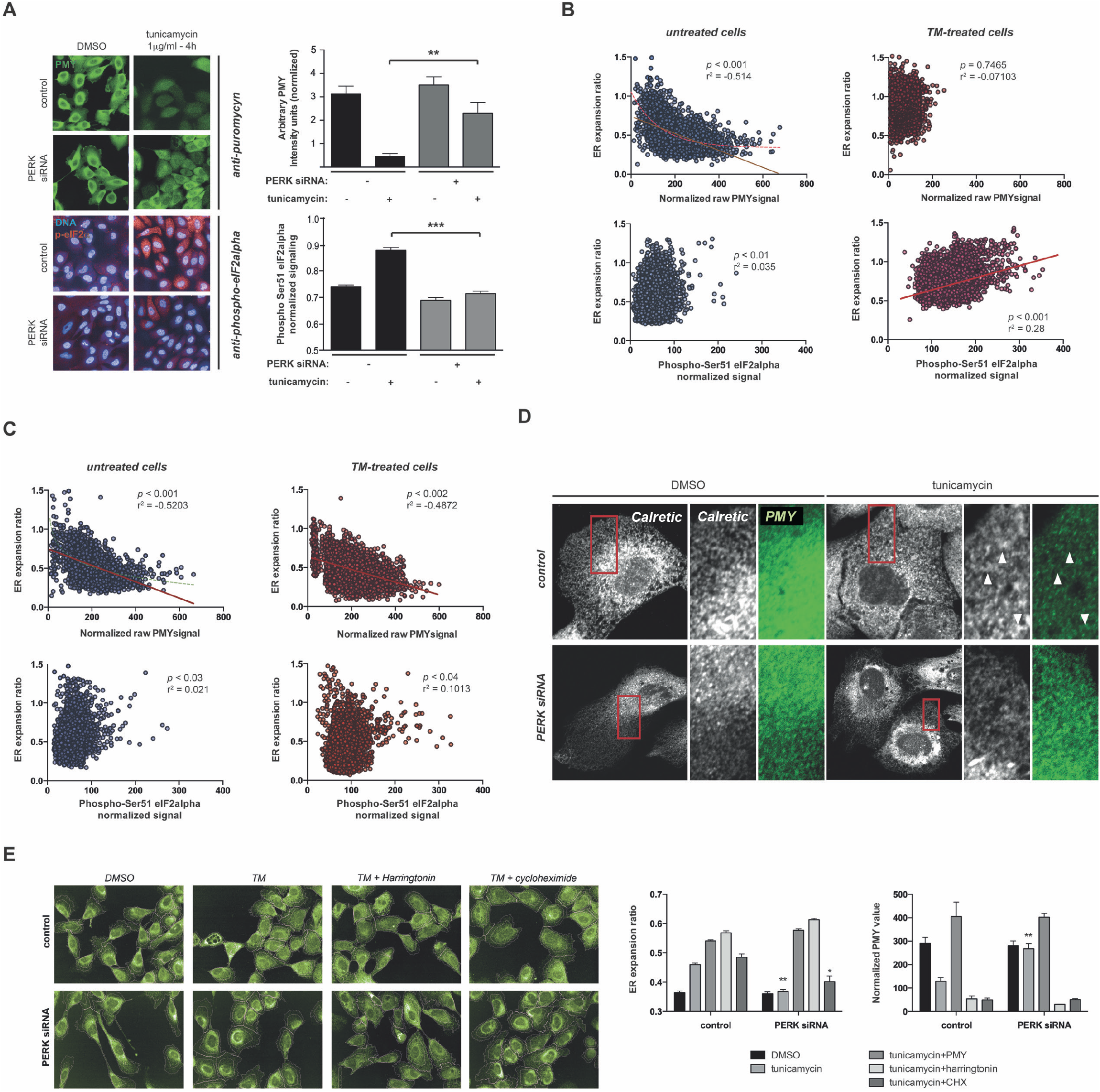
PERK-dependent translation initiation inhibition correlates with peripheral ER remodeling. **(A)** Puromycylation assays reveal a correlation between PERK-dependent protein translation shutdown and eIF2a phosphorylation upon exposure to acute ER stress in MCF10A cells. Data are derived from eight biological replicates (~2000 cells per well) across indicated treatments for each label. **(B, C)** Plotting of values recorded for single MCF10A cells from cultures either mock-transfected [B] or PERK-depleted [C]. Note correlation and statistical significance of this correlation across treatments between labels, and how this relationship is no longer sensitive to ER stress induction in PERK-depleted cells **(D)** Single confocal planes of MCF10A cells double-stained for ER (calreticulin, white) and active translation (PMY, green) across indicated siRNA and compound treatments. Note that peripheral sheet-like patches in mock-transfected cells exposed to ER stress are particularly devoid of PMY label (arrowheads). **(E)** Pharmacological inhibition of protein translation initiation, but not translation elongation, reverts the impairment for peripheral ER remodeling in PERK-depleted cells. PMY stands for both puromycylation label, and sustained inhibitory treatment. CHX: cycloheximide. Data is derived from four biological replicates (~2000 cells per well) across indicated treatments. Statistical significance across assays was assessed by paired t-Student test; *: p <0.05; **: p <0.01; ***: p <0.005. n.s.: *p* >0.05

Importantly, knockdown of PERK blunted these responses (fig. 3C). Detailed confocal analysis showed that peripheral expanded ER structures in cells subject to ER stress showed reduced overlap with PMY signal, supporting that translational shutdown is a required step for ER adaptive remodeling (fig. 3D). Of note, induction of Fv2E-PERK oligomerization was associated with significant increases in the relative expansion of the ER, which correlated with eIF2alpha phosphorylation and translation attenuation levels, in the absence of ER stress (fig. S3C). Thus ER expansion during stress correlates with reduced polysome assembly and suppression of translation.

We next tested whether bypassing PERK-dependent shutdown of translation initiation using different inhibitors of protein synthesis, could rescue the suppression of ER expansion associated with PERK depletion/inhibition during ER stress challenge. Brief exposure of tunicamycin-challenged cells to translation initiation inhibition (harringtonin or puromycin) completely reverted the ER collapse specifically associated with abolition of PERK function (fig. 3E). Interestingly, the rescue was associated rather specifically with blockade of translation initiation and polysome assembly, because exposure to cycloheximide, a drug intervening the elongation step of protein translation and stabilizing polysomes in short term timeframes, was unable to revert the PERK siRNA phenotype (fig. 3E), even under conditions that profoundly inhibit *de novo* protein synthesis^29^. Our observations support a working model whereby deficient regulation of translation initiation *per se*, and not potential client protein overload, is the cause leading to blunted ER redistribution in PERK-deficient cells during ER stress.

We performed a set of experiments to elucidate the requirement for eIF2α we had observed upon disruption of PERK activity. Cotransfection of an eIF2α S51D phospho-mimetic mutant, but not of a wild type construct, rescued the ER collapse observed in PERK-depleted, tunicamycin-challenged cells (fig. 4A-C). Notably, we observed moderate but significant increases of ER relative expansion in the absence of ER stress challenge both in wild-type and PERK-depleted cells upon ectopic expression of the phospho-mimicking mutant (fig. 4B and C), supporting a model whereby active mRNA translation *per se* is the main factor dictating relative ER distribution and architecture in our experimental model.

**Figure 4.**
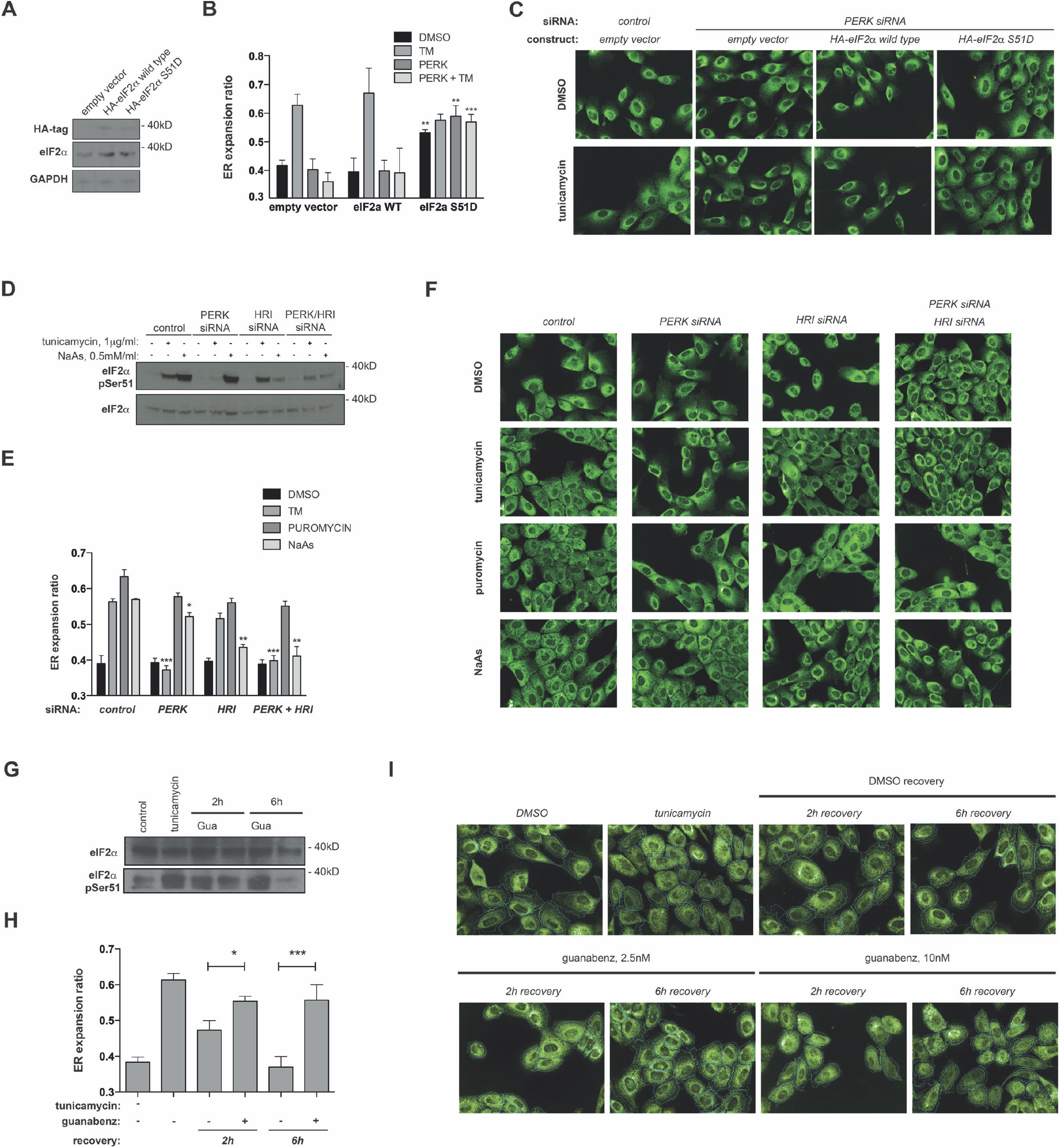
eIF2a phosphorylation is required for PERK-dependent induction of peripheral ER remodeling during ER stress. **(A-C)** Cells stably expressing a phosphomimicking eIF2a-S51D or the wild type counterpart [A, westernblot analysis] were assessed for peripheral ER remodelling across indicated conditions [B,C]. Data is derived from four biological replicates (~2000 cells per well). **(D-F)** Cells transfected with indicated siRNA duplexes were exposed to indicated treatments, and analyzed for eIF2a phosphorylation [D] or peripheral ER expansion [E, F]. Data is derived from four biological replicates (~2000 cells per well). **(G-I)** MCF10A cells were exposed to tunicamycin for 4h, and then fixed, are allowed to recover for indicated times in the presence or absence of guanabenz. Whole-cell extracts were subjected to western blot analysis for indicated markers [G]; fixed samples were processed for immunostaining for calreticulin and analyzed for relative peripheral ER content. Data is derived from four biological replicates (~2000 cells per well). Statistical significance across assays was assessed by paired t-Student test; *: p <0.05; **: p <0.01; ***: *p* <0.005. n.s.: *p* >0.05

To further characterize the requirement of eIF2α phosphophorylation for ER expansion during ER stress, we performed experiments where we stimulated the activity of an alternative eIF2α kinase during ER stress in PERK-deficient cells. Sodium arsenite specifically stimulates eIF2α phosphorylation through HRI/EIF2AK1 (fig. 4D). Of note, exposure to sodium arsenite led to ER expansion in the absence of ER-targeted drugs, both in wild type cells and PERK-depleted cells (fig. 4E and F).Moreover, sodium arsenite rescued the ER collapse phenotype associated with PERK depletion/inhibition (fig. 4E and F). In accordance with the idea that ER expansion is dependent on eIF2α phosphorylation status, depletion of the HRI kinase prevented sodium arsenite-derived rescue (fig 4E and F). Finally, we tested the effect of artificially delaying the dynamics of eIF2alpha phosphorylation using a specific inhibitor of the PPP1R15B phosphatase, guanabenz^38^; which blocks eIF2alpha dephosphorylation upon clearance of ER stress (fig. 4G). Consistent with an intrinsic role for eIF2alpha-dependent regulation of translational activity in the cell to determine ER subcellular distribution, exposure to guanabenz delayed the reverse redistribution of the ER associated with ER stress clearance in wild type MCF10A cells (fig. 4G-I). Taken together, these observations support that the inhibition of translation initiation and polysome assembly through eIF2alpha phosphorylation underpins PERK-mediated ER expansion.

### The integrity of non-centrosomal MTs determines PERK-dependent regulation of ER architecture

We next decided to look for additional players in this translation activity-dependent regulation of ER architecture. Using a double siRNA screening approach, we queried a focused list of structural ER “shapers”, known to play specific roles in ER subdomain definition (fig. 5A) to assess whether their inhibition suppressed or enhanced the ER morphogenesis defects observed upon PERK depletion. siRNAs targeting these proteins were effective and did not affect the levels of PERK mRNA (fig. S4A). This small siRNA library was either transfected alone or cotransfected with PERK-targeting siRNA. Finally, these combinations and their corresponding control and PERK siRNA-only controls were exposed to either tunicamycin or vehicle (DMSO) alone. Depletion of REEP4, p180/RRBP1 and Climp63/CKAP4 rescued the PERK siRNA phenotype of ER ‘collapse’ upon tunicamycin exposure (fig. 5B; highlighted). But this rescue did not correlate with an alteration of protein translation, as inferred from PMY staining (fig. S4B). Curiously, these hits are mostly conserved in higher metazoans, and orthologs are not found across clades where EIF2AK3/PERK is absent^39^, suggesting a co-evolved functional relationship (fig. S4C). We conclude ER collapse in PERK-deficient cells involves an altered regulation of specific ER shapers.

**Figure 5.**
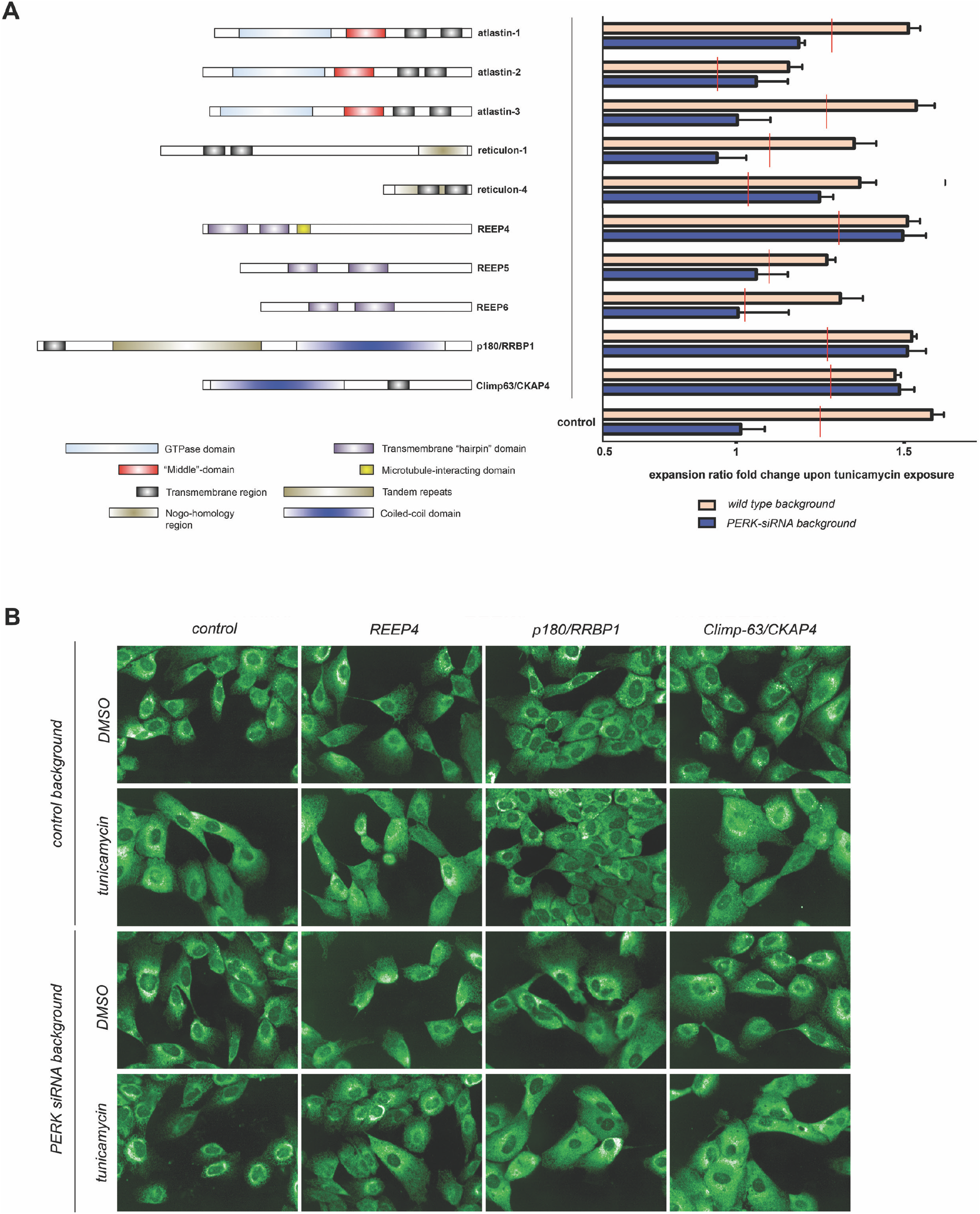
Impaired peripheral ER remodeling in PERK-depleted cells is reverted upon depleting specific ER ‘shaping’ proteins that link the ER to the microtubule cytoskeleton. **(A)** The indicated ER shapers were targeted for transient depletion in MCF10A cells by transfecting siRNA pools, either alone or in combination with siRNA duplex targeting PERK. 48h later, cell subsets were mock-treated with vehicle (DMSO) or exposed for 6h to tunicamycin, and samples were fixed and processed for calreticulin immunofluorescent staining and automated microscopy. Increase in ‘ER expansion ratio’ in tunicamycin-treated subsets, as compared to their DMSO-treated counterparts, is shown for each siRNA group. Red lines indicate the threshold of significance (*p*<0.05) for the difference between single siRNA treatment and the PERK-ER shaper double siRNA combination. Highlighted targets exhibit behavior comparable to mock-silenced cells, and rescue of the phenotype associated with PERK depletion. Data were derived from four biological replicates (~2000 cells per well). **(B)** Representative images of indicated siRNA combinations. Statistical significance across assays was assessed by paired t-Student test; *: *p* <0.05; **: *p* <0.01; ***: *p* <0.005. n.s.: *p* >0.05

Because REEP4, p180/RRBP1 and Climp63/CKAP4 are involved in the linkage between the ER and the microtubule (MT) network^25,28,40^, we tested the involvement of MT stability and organization in the phenotypes associated with PERK deficiency. Disruption of MT polymerization upon exposure to nocodazole had a profound impact on ER architecture, promoting a relative redistribution and an increase in sheet-like domains in untreated wild-type cells (fig. 6A). Nocodazole treatment also completely reverted the observed ER collapse associated with PERK depletion (fig. 6A). Importantly, this effect is not dependent on translation activity *per se*, because nocodazole-treated cells are still competent in engagement of ribosomes in protein synthesis at the times and concentrations tested (fig. S5A). Thus, we conclude ER collapse in the absence of PERK activity is driven by interactions between ER shapers and MTs.

**Figure 6.**
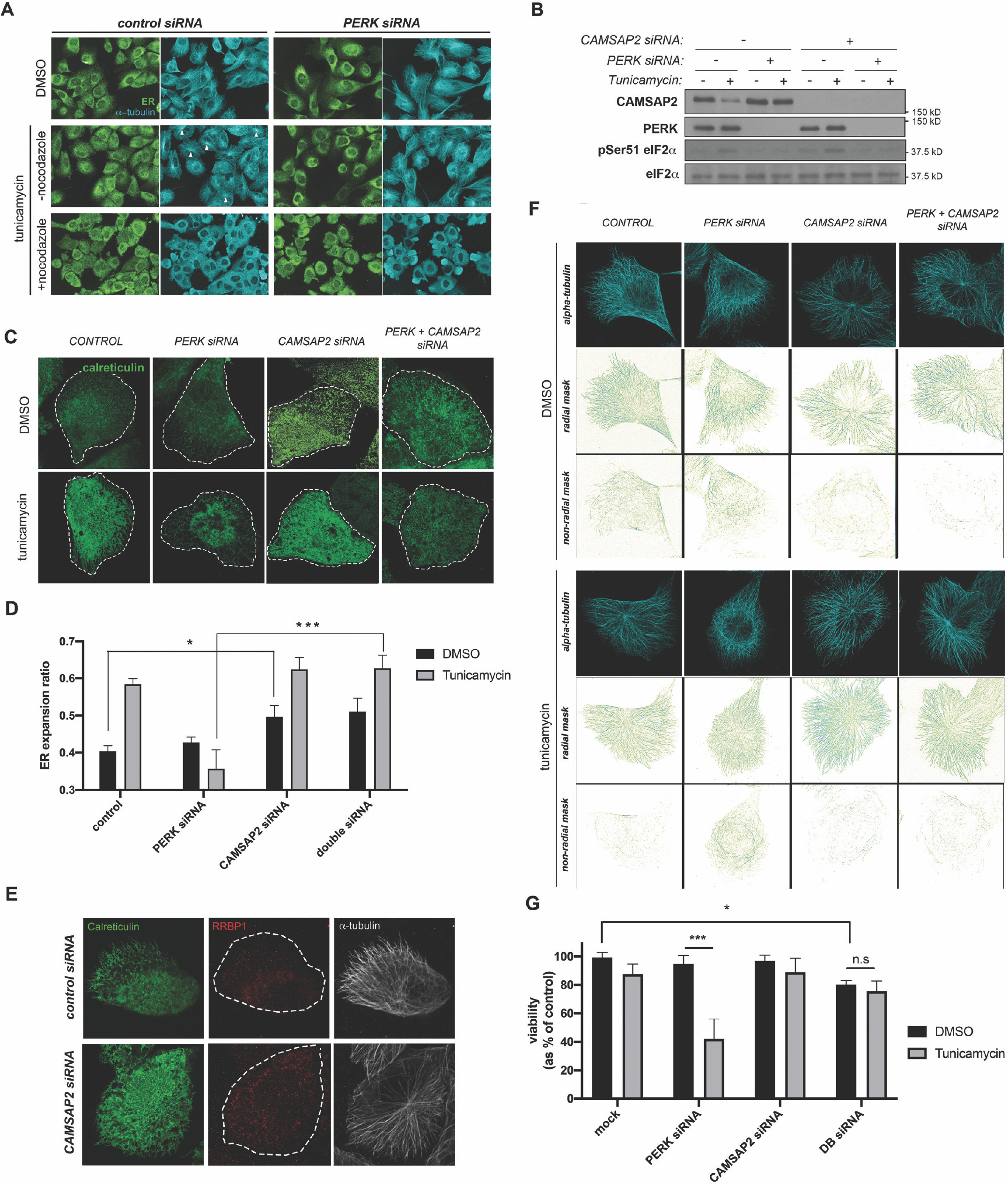
Non-centrosomal microtubules are specifically required for PERK depletion to induce a phenotype of impaired peripheral ER remodeling. **(A)** MCF10A cells were subjected to indicated siRNA and small compound treatments, fixed and immunostained for ER and microtubules, and imaged. Arrowheads indicate cells with clear radial disposition of microtubules. **(B-D)** MCF10As were reverse-transfected with indicated siRNA combinations and exposed to indicated small-compound treatments, and analyzed by western blot [B] or quantitative imaging [C, D]. Data in [D] were derived from four biological replicates (~2000 cells per well). **(E)** Triple immunostaining for endogenous calreticulin (green), RRBP/p180 (red) and α-tubulin (grayscale) in wild type MCF10A cells and cells depleted for CAMSAP2; cell boundary is highlighted with dashed lines. **(F)** Analysis of radial and non-radial microtubule signals from MCF10A cells subjected to indicated siRNA and small-compound treatments. **(G)** MTT viability assay across indicated conditions. Data were derived from eight biological replicates. Statistical significance across assays was assessed by paired t-Student test; *: *p* <0.05; **: *p* <0.01; ***: *p* <0.005. n.s.: *p* >0.05

We wondered whether specific MT populations might be involved in linker morphogenesis. Two major classes of microtubule populations exist: microtubules nucleated at centrosomes, and non-centrosomal microtubule bundles^41^. We first assessed whether centrosomal microtubules were required to determine ER cell distribution and architecture by depleting centrosome structures from MCF10A cells using the polo-like kinase 4 (PLK4) inhibitor centrinone^42^. The ER of cells devoid of centrosomal MTs were phenotypically similar to those in PERK/EIF2AK3 knock-down cells, indicating that depletion of MTs nucleated at the centrosome does not rescue the ER expansion defects associated with PERK inhibition (fig. S5B). Next, we depleted mRNAs encoding the calmodulin regulated spectrin associated protein family member 2 (CAMSAP2), a well-established minus-end stabilizer of non-centrosomal MTs^41,43^ (fig. 6B). Cells deficient for stabilization of non-centrosomal MTs alone exhibited a moderate phenotype of increased peripheral ER cistern-like structures (fig. 6C and D). Importantly, CAMSAP2-deficient cells simultaneously knocked down for PERK rescued the collapsed perinuclear ER and impaired ER redistribution associated with PERK RNAi alone (fig. 6C and D). In accordance with increased peripheral extension of sheet-like structures, CAMSAP2-depleted cells showed higher signal density for RRBP1 immunostaining in the cell periphery (fig. 6E). It must be noted that neither of these interventions affected significantly overall translation activity (fig. S5C), suggesting that they do not provoke unspecific reduction of ER polysome density. The most parsimonious explanation for these observations is that PERK deficiency leads to a collapsed ER phenotype by stabilizing translation-dependent interactions between ER shapers and non-centrosomal pools of MTs.

We wondered whether ER stress and PERK activity might also affect specifically MTs organization directly We appreciated in previous experiments that acute ER stress induction was associated with the presence of prominent discrete MT nucleation centers in wild-type cells, whereas PERK-depleted cells did not exhibit such features (see fig. 6A, white arrowheads). To study this phenomenon in higher detail, we obtained MT image sets using STED microscopy from cells exposed to acute ER stress, either transfected with scrambled siRNA, or depleted for PERK, CAMSAP2, or both proteins simultaneously (fig. 6F; see also fig. 6B). As expected, depletion of CAMSAP2 alone led to a relative decrease in non-radial MT structures in the presence of vehicle indicating a loss of non-centrosomal MTs(DMSO). Wild-type cells exhibited also decrease in non-radial MT structures when exposed to acute ER stress, suggesting ER stress suppress non-centrosomal MT polymerization. Of note, this phenotype correlated with a significant decrease in CAMSAP2 total protein of c. 50%, which was not observed in PERK-depleted cells (see fig. 6B, lanes 2 vs 4). However, such decrease in non-centrosomal MTs was not observed cells depleted for PERK (fig. 6F). These observations suggest that PERK suppresses non-centrosomal MT polymerization. Finally, cells depleted of both PERK and CAMSAP2 had very few non-centrosomal MTs suggesting PERK ability to suppress non-centrosomal MT polymerization is upstream of CAMSAP2. Of note, CAMSAP2 depletion attenuated tunicamycin-induced cell viability reduction in PERK-deficient cells, albeit viability in DMSO-treated double-knockdown cells was also slightly affected (fig. 6G). These observations support that non-centrosomal microtubules are suppressed in a PERK-dependent fashion during ER stress to promote ER expansion. Taken together with previous observations, this suggests that active polysomes allow for the anchoring of ER shapers to non-centrosomal MTs in PERK-deficient cells, leading to a collapsed ER structure upon ER stress induction.

### Disruption of PERK kinase activity impacts cell protrusiveness and motility

Non-centrosomal MT arrays can regulate cell protrusiveness and migration^41,43^. Of note, and as suggested by our initial screening image datasets, depletion of PERK kinase in cells led to increased protrusion size but reduced protrusion number, and increased polarity, as compared to wild-type cells (Fig. 7A-C). These observations are in agreement with an interpretation that PERK activity could regulate cell polarity by suppressing the polymerization of non-centrosomal MTs, which contribute to cell protrusiveness^43^. In support of this idea, increased protrusiveness in PERK deficient cells was not observed in PERK deficient cells also depleted for CAMSAP2 (Fig. 7A-C).

**Figure 7.**
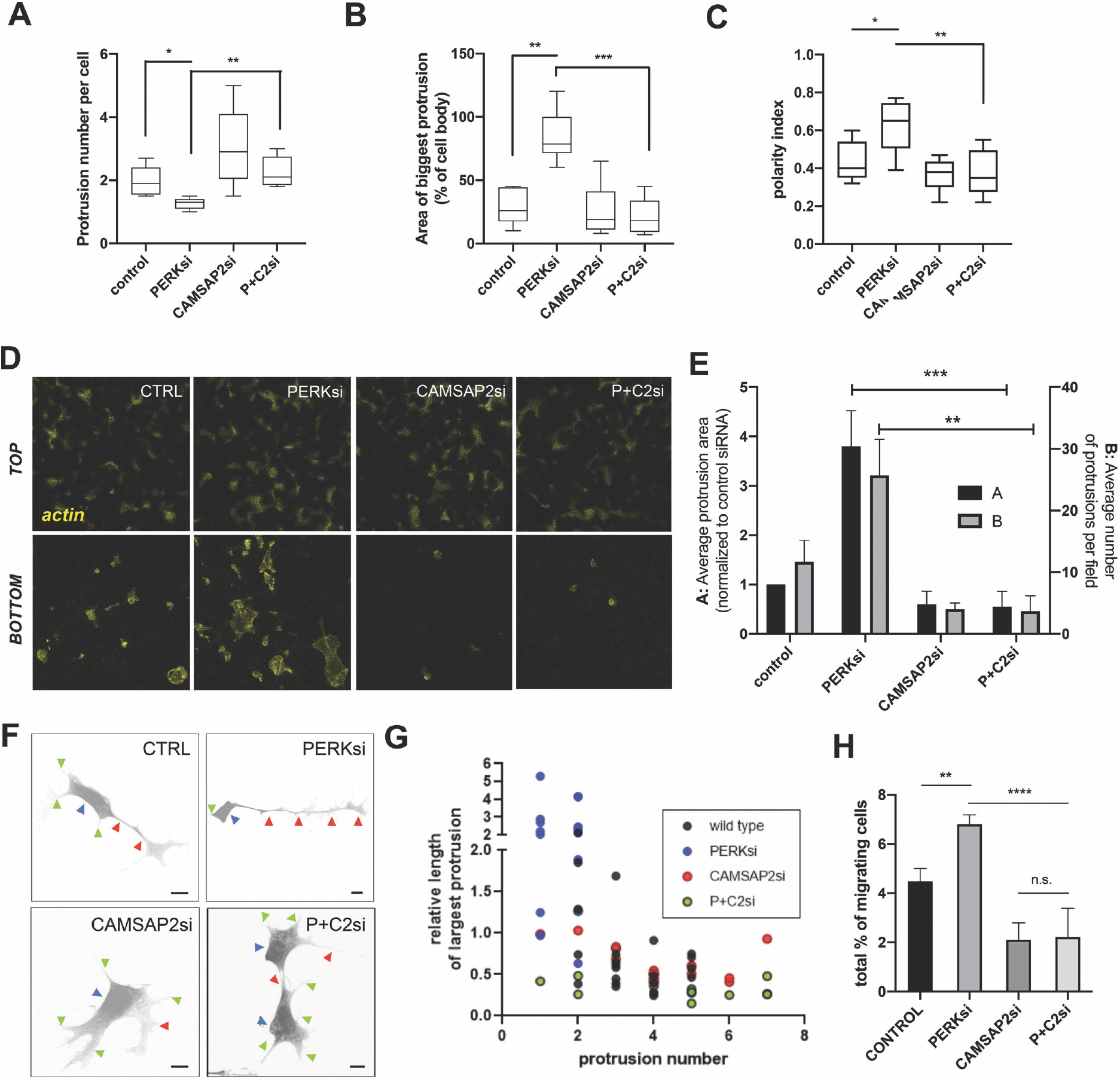
PERK depletion impacts on cell protrusiveness and polarized migration in a manner dependent on non-centrosomal microtubules. **(A-C)** Features indicated were extracted from a minimum 25 microtubule- and Golgi immunostained, cytoplasm-counterstained cell confocal images across indicated conditions. **(D-E)** MCF10As subjected to indicated siRNA treatments were allowed to protrude through Transwell™ pore membranes for 4h, then fixed and stained for actin and imaged. [E] Indicated parameters were computed from 10 independent fields per condition, from 3 biological replicates. **(F-G)** SH-Sy5y neuroblasts transfected with indicated esiRNAs were stimulated for neural differentiation (see M&Ms). Cell soma (blue arrowheads), the longest protrusion (red arrowheads) and secondary protrusions (green arrowheads) are highlighted. Scale bar: 50μm [G] The number of protrusions per cell, and the length of the longest protrusion relative to the perimeter of the cell soma are plotted for each condition (n=20 from 2 biological replicates). (**H)** MCF10As subjected to indicated siRNA treatments were allowed to migrate through 3D collagen matrices, and imaged at plate bottom, and three 50micron-apart optical sections. % of cells migrated from total cell count is indicated. Data were derived from six biological replicates. Statistical significance across assays was assessed by paired t-Student test; *: *p* <0.05; **: *p* <0.01; ***: *p* <0.005. n.s.: *p* >0.05. Error bars depict S.E.M.

To further assess the apparent directional protrusiveness phenotype of PERK-depleted cells, we performed Transwell™ migration assays. We observed a higher degree of protrusions passing through membrane pores when either depleting cells for the PERK kinase using siRNA, as well as upon exposure to the PERK allosteric kinase inhibitor (fig. 7D, see also fig. S6A). Importantly, this increase in protrusiveness was abrogated by simultaneous depletion of the non-centrosomal minus-end stabilizer CAMSAP2 (fig. 7D and E). Increased polarized protrusiveness associated with PERK depletion was also observed during neurite formation in neuroblasts (fig. 7): PERK-deficient cells exhibited very long unique protrusions, as compared to wild type cells. Again, simultaneous depletion of CAMSAP2 reverted these phenotypes.

We sought to study the correlation of these phenotypes with cell motility. We assessed the impact of PERK signaling disruption on 3D cell migration. PERK inhibition enhanced cell migration through a 3D collagen matrix, and this phenotype was inhibited upon depletion of the non-centrosomal MT stabilizer CAMSAP2 (fig 7F). In accordance, cells plated on soft collagen matrices exhibited long protrusions (fig. S6B). Of note, depletion of the ER-MT linker RRBP1/p180 also reduced the increase in elongated protrusions and abrogated the increase in 3D cell migration (fig. S6B and C). Artificial PERK activation in the synthetic Fv2E-PEK model by exposure to the homodimerizer AP20178 had an opposite effect on 3D cell migration, further supporting a role for PERK-dependent signaling on the regulation of cell motility (fig. S6D). We propose a model whereby ER-MT reciprocal regulation through PERK-dependent translation control has a dual impact on both regulated ER remodeling, and non-centrosomal MT-dependent cell polarity and migration.

## DISCUSSION

Eukaryotes evolved complex functional programs that continuously monitor ER physical integrity and function and coordinate different responses in the cell to primarily adapt its function, collectively termed Unfolded Protein Response (UPR). Here, we contribute evidence that PERK-dependent control of protein translation modulates the reciprocal regulation of ER and non-centrosomal microtubules, to both drive ER remodeling during acute ER stress, and regulate non-centrosomal MT-dependent cell protrusiveness and polarity.

Both ribosome association and microtubule cytoskeleton integrity can affect ER architecture, as well as the localization of ER shapers^29^. Here, we demonstrate that this relationship is controlled by the PERK kinase through eIF2a phosphorylation-mediated translation shutdown. ER architecture is tightly regulated by different ‘shaper’ proteins, capable of defining different ER membrane domains^2^. The targeting of ER shaping proteins must be controlled to allow for the dynamic remodeling of these membrane structures for cell adaptation to different functional states, and different mechanisms have been suggested, including posttranslational modifications and discrimination of differential membrane curvature^28,44–46^. While a detailed picture of the precise mechanisms at play requires further investigation, our results support that the control of the dynamics of ternary complexes involving polysomes, microtubules and specific ER-anchored proteins, is pivotal. The fact that depletion of each of the ER-MT linkers we have found in our study (p180, Climp63 and REEP4) has a significant impact on the ‘collapsed’ phenotype we observe in PERK-deficient cells suggests that each of these components is necessary, but not sufficient, for the coupling of ER and MTs in epithelial cells. Given the impact of these ER-MT tethers on MT network architecture (and particularly, on the non-radial bundling of MTs when dysregulated)^25,28^, it will be interesting to also explore the specific role of each of these proteins on the dynamic stability of non-centrosomal MTs. An additional pending question is the potential interplay of these mechanisms with the reported crosstalk between the PERK-eIF2a axis and the actin cytoskeleton^47–49^; because both cytoskeletons engage in mutual regulation^50^, it is plausible to assume their crosstalk is relevant in the control of ER-MT coordination we report here.

Physical ER expansion and shape remodeling is a relevant adaptive event in cells with compromised ER function (i.e. ER stress)^4^, and closely correlates with other adaptive responses such as upregulation of protein maturation machineries^51^. PERK-dependent coupling of attenuation of client protein load and ER expansion and remodeling would ensure matching of both adaptive ER volume increase and protective translation attenuation. Moreover, appropriate expansion of ER membrane might benefit from this coupling, ensuring a limited density of ER membrane-inserted proteins. At present, despite the links with UPR signaling documented, we do not know what the relationship of our findings with mechanisms driving ER membrane biogenesis is^4,52^. We have observed that depletion of non-centrosomal microtubules attenuates the relative sensitivity of PERK-deficient cells to acute ER stress, although CAMSAP2 depletion *per se* seems to have an impact on ER homeostasis (see fig. 6). While the expansion of peripheral ER sheets is an integral aspect of cell adaptation to ER stress, future studies will be required to characterize in detail how architectural changes in the ER impact different outputs of ER function. Additional pending questions for future exploration pertain to the precise molecular mechanisms by which non-centrosomal MTs are regulated during ER stress (figure 6). CAMSAP2 levels are indeed sensitive to acute ER stress induction^53^. Apart from tight eIF2a-dependent regulation of translation, other mechanisms coordinated by PERK activation, such as autophagy^54^, may be at play

Available literature describes a relationship between ER architecture and MT dynamics, and previous studies already suggested a specialization of ER-associated MT pools^23^. What is the significance for non-centrosomal MTs to determine ER architecture? Functional segregation between centrosomal and non-centrosomal MTs is not fully understood, but an emerging major role of non-centrosomal microtubules, as suggested by studies manipulating this MT subpool, is the onset and maintenance of cell polarity and protrusiveness across cell types^41,43^. Importantly, cell polarity and protrusiveness are linked to organelle dynamics and trafficking through mechanisms that rely on the control of organelle architecture, and couple them to *de novo* membrane synthesis^55,56^. It is intuitive that their coupling to ER stress surveillance is convenient to ensure cell homeostasis across functional states.

What is the physiopathological relevance of this dynamic link between ER and non-centrosomal microtubules? Neuron differentiation and the control of neurite outgrowth and axon stabilization is largely determined by the fine regulation of ER architecture, MT organization and their reciprocal crosstalk therein^57–59^, and challenging ER homeostasis can compromise this specialized form of cell protrusion stabilization^60^. Moreover, non-centrosomal MTs appear to be specifically involved in these mechanisms^57,61,62^. Because eIF2a phosphorylation and PERK activity modulate memory stability and learning^63–65^, it will be interesting to study the relevance of our findings in these contexts. Future studies focusing on the impact of these mechanisms on directional vesicle trafficking and *de novo* membrane formation, which seem to be relevant for protrusion formation, may also shed light on these questions^56^. We have studied invasiveness phenotypes derived from PERK depletion in mammary epithelial cells, as related to increased polarity. PERK activity has been studied in the context of mammary acini development and tumorigenesis, and its complex impact on tumor cell survival and tissue organization^35,66–68^. Interestingly, recent studies support a role for CAMSAP2 in tumor cell invasiveness^69^. The emerging picture is complex, because the impact of intervening PERK signaling on tumor cells is highly contextual and simultaneously affects different aspects of tumor cell biology: cell survival and adaptation to nutrient deprivation and adverse environment, adhesion signaling and accommodation of altered secretory phenotypes. Our results contribute a novel additional perspective to this complex picture and suggest novel opportunities for synergistic intervention of tumor cell biology by shutting down PERK-dependent prosurvival signaling (autophagy, ROS management) and cell invasiveness.

## MATERIALS AND METHODS

### Cell culture, transfection and reagents

All cell maintenance and experiments were performed in a standard humidified incubator at 37°C and 5% CO_2_, unless otherwise stated. Low-passage MCF10A cells, RasV12-transformed MCF10A ATI cells, and MCF10A-derived stable cell lines were cultured in DMEM-F12 Glutamax® medium (Gibco) supplemented, unless otherwise stated, with 5% heat-inactivated horse serum (Sigma), 1μg/ml bovine insulin, 1μM hydrocortisone, 50U cholera toxin, and 100ng/ml epidermal growth factor (EGF)-selection (puromycin, 5ng/ml) was applied in stable cell lines. HeLa, Sh-Sy5y and MDA-MB231 cells were grown in high glucose DMEM supplemented with 10% heat inactivated fetal bovine serum (Gibco).

MCF10A and MCF10A-ATI cells were a kind gift from Claire Isacke (ICR, UK); MCF10A/Fv2E-PERK and MCF10A/PERKΔC stable cell lines were generously provided by Julio Aguirre-Ghiso (Mount Sinai Hospital, USA). ER bulk content was analyzed by flow cytometry after brief pulse-labeling with BODIPY FL-ER tracker (Molecular Probes, ThermoScientific) freshly resuspended cells after indicated treatments and analyzed on a LSR Fortessa station. For siRNA reverse transfection, Lipofectamine RNAiMAX (LifeSciences) reagent was used following supplier’s recommendations. Plasmid transfections were performed using Lipofectamine 3000 (LifeSciences) for 24h. Tunicamycin, puromycin, sodium arsenite, cicloheximide, nocodazole, guanabenz and Hoescht 33258 were obtained from Sigma. Thapsigargin was obtained from LifeSciences-Invitrogen. The GSK202646 PERK inhibitor and harringtonin were purchased from Tocris. AP20187 homodimerizer was purchased from Selleckchem, centrinone was purchased from R&D systems. Methanol-free, 16% paraformaldehyde-PBS was purchased from ThermoScientific. Secondary antibodies and fluorescent conjugates were purchased from Molecular Probes. siRNAs were obtained from Dharmacon, esiRNAs were purchased from Sigma. A table listing primary antibodies and siRNA/esiRNAs used is available as supplementary table 1.

### cDNA constructs, establishment of stable cell lines, and transient transfections

MCF10A-EGFP-Sec61β cell line was obtained by lentiviral transduction (MOI: 5; Viral Vector unit, CNIC, Spain; lentiviral vector cloned on pRRL-IRES-mCherry from NdeI/MluI digested parent AcGFP-Sec61B vector Addgene #15108) and two subsequent rounds of stringent sorting (FACS Aria, BD Biosciences) of EGFP positive cells with a fluorescence range within one order. Reverse transfection proceedings for transient silencing have been detailed elsewhere. For neurite extension assays, SH-Sy5y cells lentivirally transduced with a pRRL-IRES-EGFP vector (MOI: 1; Viral Vector unit, CNIC, Spain) were reverse transfected with indicated esiRNAs and incubated for 48h, then switched to differentiation medium (1.5% FBS, 1μg/ml bovine insulin, 1μM hydrocortisone, 10μM *trans*-retinoic acid (Sigma®), and 5U brain-derived neurotropic factor (BDNF, Sigma®)) and incubated for a further 48h before being processed for immunofluorescence. eIF2a WT and S51D constructs were cloned on pcDNA3.1-HA from Addgene constructs #21807 and #21809. In experiments detailed in figure 4A, where simultaneous transfection of cDNA constructs was required, cells were reverse transfected as detailed above, and 24h after directly transfected using the Lipofectamine 3000 reagent.

### High-throughput assays, immunostaining and acquisition

For high content imaging, cells were reverse transfected using Lipofectamine RNAiMAX reagent, and 40ng of siRNA, on optical CellCarrier 384-well plates, on a final volume of 40μl. Liquid handling, fixation and immunostaining was performed using a robotic station (HCS Explorer, PerkinElmer) according to previously detailed protocols^70^. Acquisition and automated image analysis was performed with an Opera HCS II spinning disk confocal microscope.

### Superesolution confocal microscopy and transmission electron microscopy (TEM)

A Leica SP8 3X STED spectral confocal microscope, equipped with hybrid photomultipliers, a CCD digital camera and a STED station with two separate depletion lasers (592 and 660nm) was used, with a 100X/1.4NA immersion objective. Sequential (between stacks) acquisition used 100% power for STED depletion and 70% power for illumination. Samples were prepared according to the manufacturer’s recommendations.

For TEM, cells grown on 100-mm dishes treated as indicated were fixed with 4% paraformaldehyde and 2% glutaraldehyde for 120 min at room temperature. Upon gentle scrapping, postfixation was carried out with 1% OsO4 and 1.0% K3Fe(CN)6 in H2O at 4 °C for 60 min. Samples were dehydrated with ethanol and embedded in Epoxy, TAAB 812 Resin (TAAB Laboratories) according to standard procedures. Ultrathin (80 nm) sections were stained with saturated uranyl acetate and lead citrate and visualized with a JEOL JEM 1010 (Tokyo, Japan) electron microscope at 80 kV. 16-bit images were recorded with a 4 k × 4 k CMOS F416 camera from TVIPS (Gauting, Germany), typically at 12000X magnification.

### Image analysis

Image analysis proceedings are detailed together with full script file as supplementary data, and comprehended the following steps: (1) nuclei and cell segmentation; (2) filtering of artifacts and out-of-focus objects; (3) definition of subcytoplasmic regions, and (4) gathering of ER-related features. ImageJ analysis pipelines used in experiments shown in figures 6 and 7 have been previously reported^43^.

### Protein analysis

Cell lysates were harvested in colorless, non-reducing sample buffer (2% SDS, 150KCl, 20mM Tris pH6.8, 10% glycerol, protease and phosphatase inhibitors), quantitated through Bradford assay and normalized, and supplemented with DTT and bromophenol blue to a final concentration of 0.2mM and 0.012%, respectively. ~10μg/sample were loaded on 10-12.5% SDS-polyacrylamide gels. After electrophoretic separation and blotting in conventional non-SDS, 20% methanol conditions to PVDF membranes, samples were probed with the indicated antibodies according to standard protocols^31^. Signal was developed with the ECL Plus system (PerkinElmer).

### Puromycilation assay

Our protocol was a slight modification of procedures published previously^37^. Briefly, cells to be labeled were exposed in a ~45” pulse to a low concentration of puromycin (500ng/ml) and immediately washed twice in complete medium and fixed with 4% PFA. Samples were permeabilized 15min in 0.2% triton-X100 PBS, washed twice in 0.05% TX-100 PBS, and blocked for 2h with 2% BSA in PBS. Standard immunofluorescence procedures were thereon applied to label with the 2B4 anti-puromycin monoclonal antibody, developed by A. David and coworkers and publicly available through the Developmental Studies Hybridoma Bank repository (entry ♯PMY-24B)

### Cell viability assays

Cell viability was inferred by the MTT colorimetric assay. Briefly, cells were treated with either vehicle (DMSO) or tunicamycin (5μg/ml, 36h), and then exposed to 50μg/ml MTT (3-(4,5-dimethylthiazol-2-yl)-2,5-diphenyltetrazolium bromide at 37 °C for 3 hours in the dark. Precipitated formazan was solubilized in DMSO, and absorption was measured at 542nm in a spectrophotometer. Absorption values were referred to control condition as 100%.

### Transwell migration assays

8.0 μm Ø- pore PET hanging inserts were pre-coated with collagen as recommended by the supplier (Millipore). Cells were seeded in a 500μl volume of reverse transfection mixture and allowed to migrate for ~4h. Inserts were then placed on pre-warmed 4% PFA-PBS, and processed for immunostaining following standard protocols. Images were acquired on a Zeis Axiovert confocal microscope. Basal (trans) fluorescence was related to top (cis) fluorescence from the actin channel as segmented and quantitated by ImageJ standard tools from raw .lif images.

### Culture on collagen matrices

~100μl Fibrillar dermal bovine collagen I layers (1.7mg/ml) were casted on 96-well optical ViewPlate plates (PerkinElmer), following previously detailed protocols. ~4000 (MCF10A, MCF10-ATI) or ~6500 (MDA-MB231) cells were reverse transfected and plated on top of the pre-casted collagen plugs, as described, on a total volume of 50μl of complete medium. After 48h, cells were pulse-labelled with CellTracker Orange 561 (Molecular Probes), fixed by adding 50μl of pre-warmed 16% PFA in PBS, and counterstained by adding 10ng of Hoeschtt 33342. Acquisition of focal stacks was performed on an Opera HCS II spinning disk microscope, and in-focus images were manually selected for further analysis.

### 3D Collagen invasion assays

96-well optical glass plates (PerkinElmer) were coated overnight at 37oC with an sterile 1% solution of BSA to minimize cell attachment. Cells reverse transfected for 24h were detached and embedded in a 1.7 mg/ml fibrillar dermal bovine collagen I matrix at a density of 50000cells/ml, and 150μl were plated per well. Plates were immediately spun at 300g for 5min and incubated for ~4h at 37°C to allow for collagen polymerization. 50μl completion media (4X growth medium) containing the indicated treatments was added, and cells were allowed to invade the matrix overnight. Subsequently, samples were fixed and counterstained adding 50μl of 16% PFA in PBS containing 500ng/ml of Hoeschtt 33342. Plates were then imaged using an Operetta HCS system (PerkinElmer; ICR, London, UK), acquiring three optical planes with a spacing of ~50μm. Nuclei segmentation and counting per plane was performed using the Columbus automated system (PerkinElmer).

## ACKOWLEDGEMENTS

For sharing with us valuable reagents and/or suggestions, we are grateful to professors Rachael Natrajan, Francisco Sánchez-Madrid, Julio Aguirre-Ghisso, Hesso Farhan, Françoise van der Goot and Jonathan Yewdell. We acknowledge the efficient and friendly assistance of the following core services: Flow Cytometry core facility at the ICR (London, UK), Advanced Light Microscopy at CNIC (Madrid, Spain), and Electron Microscopy at CBM-CSIC (Madrid, Spain). We are deeply grateful to the Cellomics Unit at CNIC for assistance and infrastructure, and to other members of the Bakal and the del Pozo labs for their input and advice. Amine Sadok and Faraz Mardakheh (former researchers at ICR, London, UK) provided expert advice and assistance on collagen migration and Transwell™ experiments. Where indicated, plasmid and antibody resources were obtained from the Addgene and DSHB public repositories and we are grateful for their excellent non-profit service. CB and HS have been beneficiaries of the Wellcome Trust Career Development Fellowship program. MS-A was a fellow of the COFUND-IPP programme (2014). Funding support was received from the Cancer Research UK (CRUK) Programme Foundation Award (C37275/A20146) and the Stand Up to Cancer campaign for Cancer Research UK to CB; and from the spanish Ministerio de Ciencia e Innovación (MCNU; SAF2017-83130-R and BFU2016-81912-REDC), the Comunidad Autónoma de Madrid/FEDER, Spain (ref. S2018/NMT4443; Actividades de I+D entre Grupos de Investigación en Tecnologías) and the Fundació La Marató de TV3 (385/C/2019) to MAdP. The CNIC is supported by the Instituto de Salud Carlos III (ISCIII), the Ministerio de Ciencia, Innovación y Universidades (MCNU) and the Pro CNIC Foundation, and is a Severo Ochoa Center of Excellence (SEV-2015-0505).

## AUTHOR CONTRIBUTIONS

MS-A and CB conceived and designed the study. HS provided image analysis tools for initial screen. MS-A performed experimental work and analyzed data, with assistance from FNL and GF. PP-V and MA-G acquired relevant preliminary results. MAdP provided key resources and advice. MS-A and CB wrote the paper.

## SUPPLEMENTARY FIGURES

**Supplementary Figure 1.**
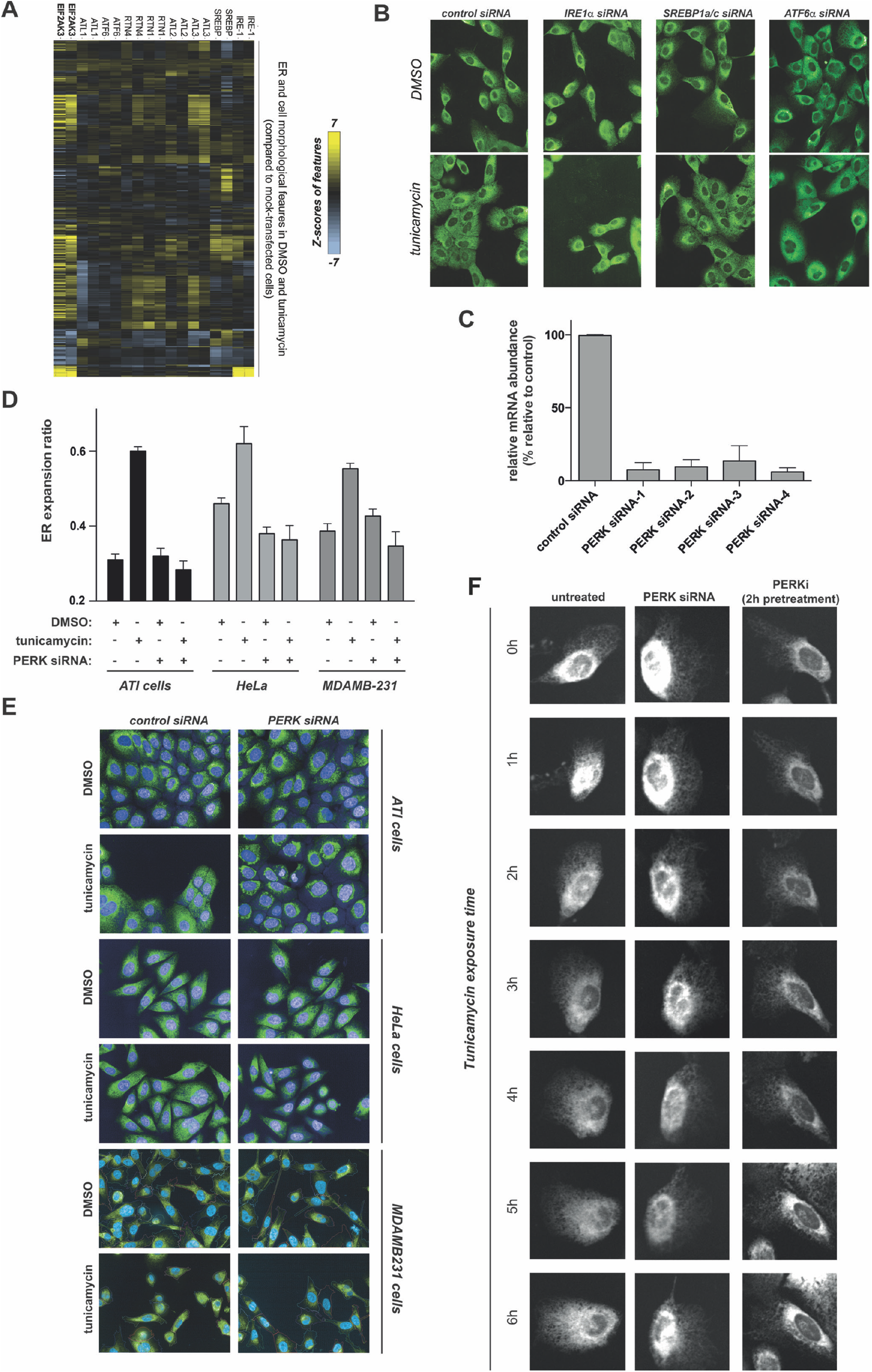
**(A)** Heatmap showcasing Z-scores extracted from 2 biological replicates (averaged for 4 replicate wells for each condition) of all 136 image features (both including treated and non-treated conditions) across indicated siRNA pairs. Hierarchical clustering renders correct pairing of siRNA duplex replicas. **(B)** Immunofluorescence images (calreticulin) of cells transfected with indicated siRNA duplexes and exposed to either vehicle (DMSO) or 1μg/ml tunicamycin for 6h. Note impaired remodelling in IRE1 and SREBP1a/c-depleted cells. **(C)** RT-PCR analysis of *EIF2AK3* mRNA levels in cells transfected with indicated siRNA duplexes (*related to figure 2A*). Data is derived from 3 technical replicates **(D, E)** Indicated cell lines were reverse transfected with an siRNA pool targeting PERK or scrambled, subjected to indicated treatments and processed for immunofuorescent staining (calreticulin; counterstained for DAPI), imaged and analysed. Graphs are derived from four independent replicates (~2000 cells per well). **(F)** Live cell imaging of MCF10A cells stably expressing an EGFP-Sec61β fusion, for indicated times and across indicated treatments. Statistical significance across assays was assessed by paired t-Student test; *: *p* <0.05; **: *p* <0.01; ***: *p* <0.005. n.s.: *p* >0.05

**Supplementary Figure 2.**
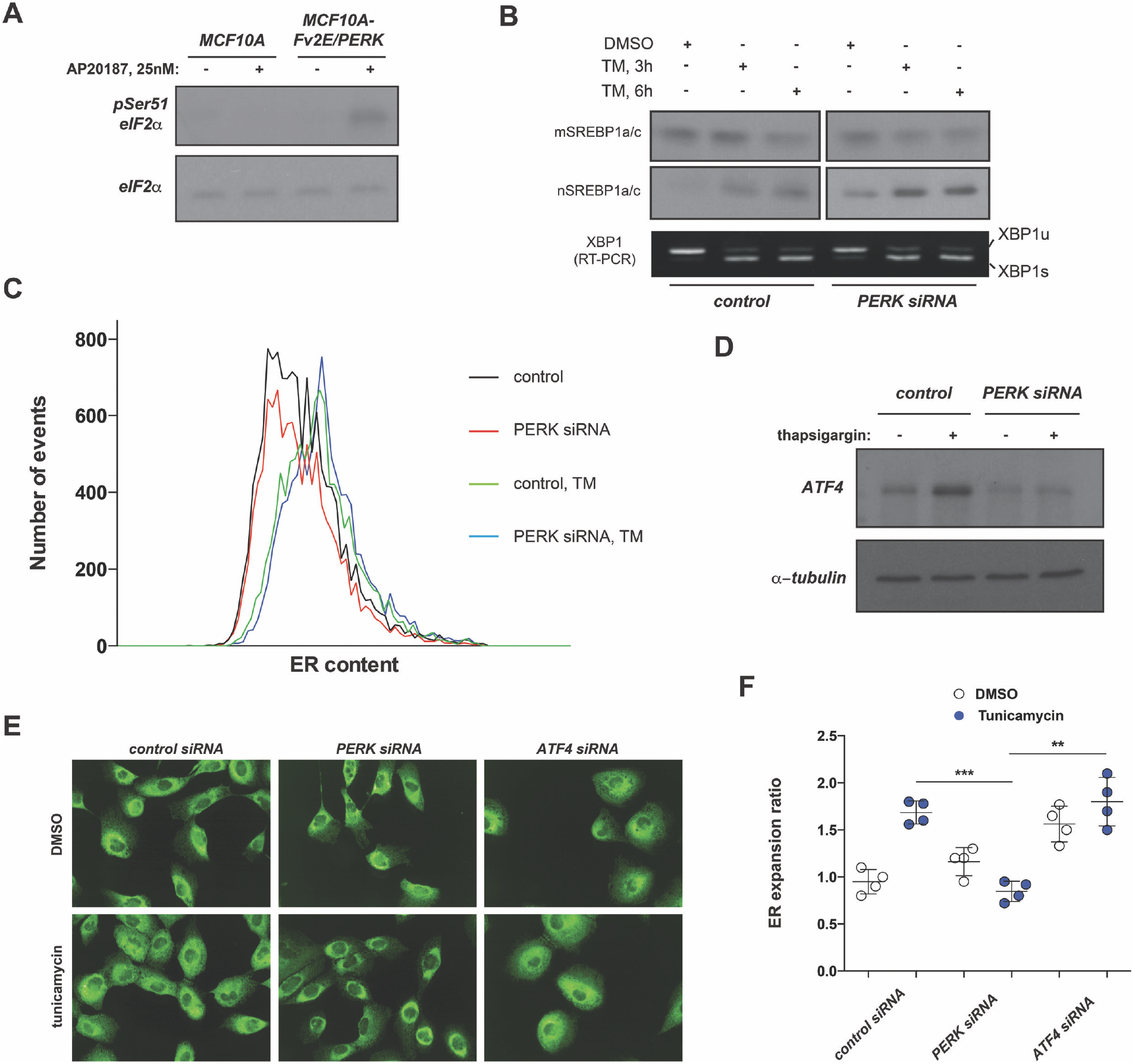
**(A)** Whole-cell extracts from MCF10A or MCF10A-Fv2E-PERK cells treated as indicated, were analyzed by western blotting with indicated primary antibodies. **(B)** Protein and total RNA was extracted from cells treated as indicated and analyzed by western blot and RT-PCR. **(C)** Cells treated as indicated were detached and labeled with ER Tracker-BODIPY FL for 15min, and analyzed by flow cytometry for ER total content. **(D)** Whole-cell extracts from MCF10A cells treated as indicated, were analyzed by western blotting with indicated primary antibodies. **(E, F)** MCF10A cells were reverse transfected with an siRNA pool targeting ATF4 or scrambled, subjected to indicated treatments and processed for immunofuorescent staining (calreticulin; counterstained for DAPI), imaged and analysed. Graphs are derived from four independent replicates (~2000 cells per well). Statistical significance across assays was assessed by paired t-Student test; *: *p* <0.05; **: *p* <0.01; ***: *p* <0.005. n.s.: *p* >0.05

**Supplementary Figure 3.**
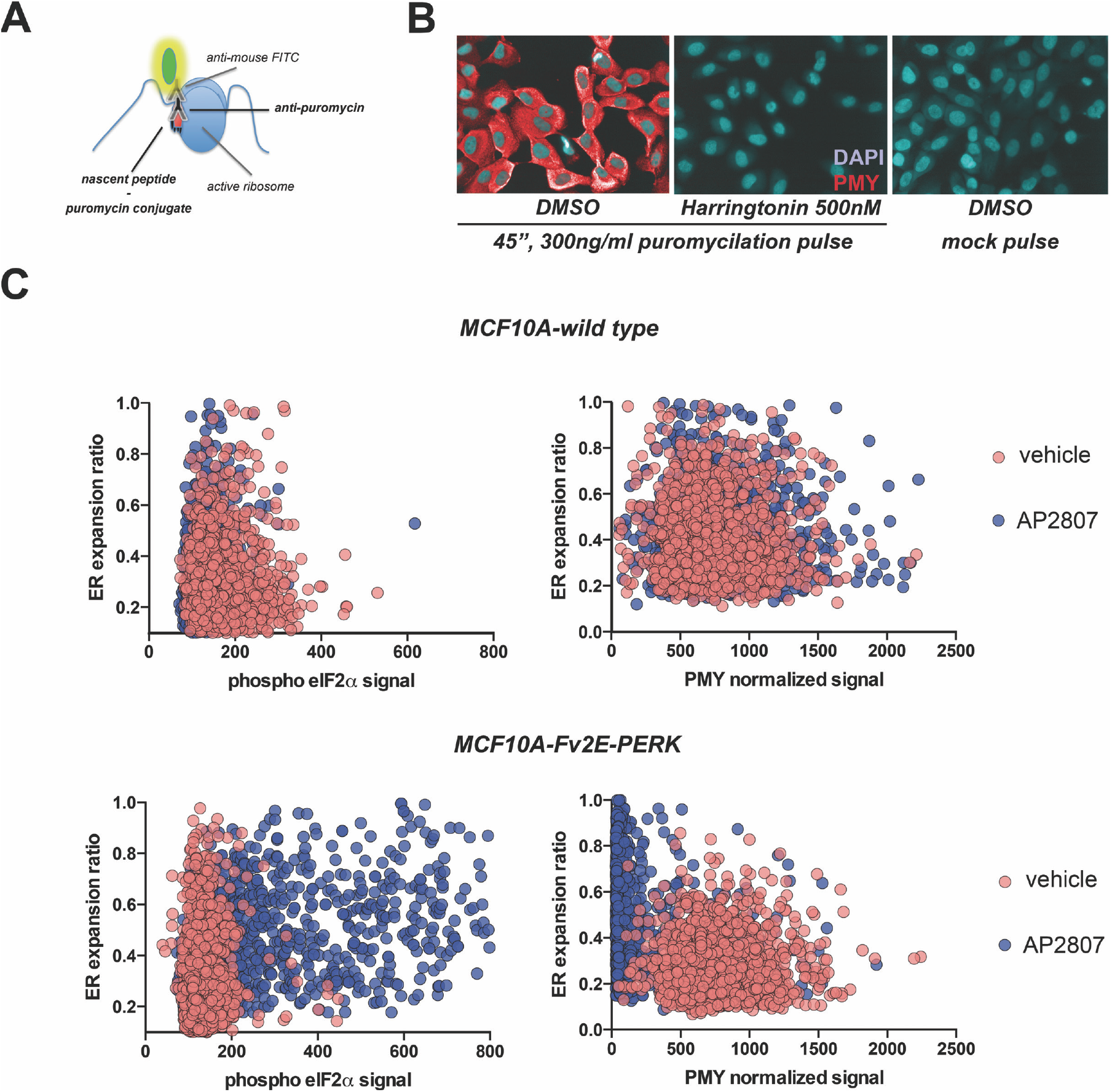
**(A)** Diagram of puromycylation immunostaining technique **(B)** Control assay showing specificity of the technique. **(C)** The synthetic Fv2E-PERK homodimerization system bypassing ER stress activation recapitulates the observations on ER expansion correlation and translation shutdown at single-cell level (related to figure 3B and C).

**Supplementary Figure 4.**
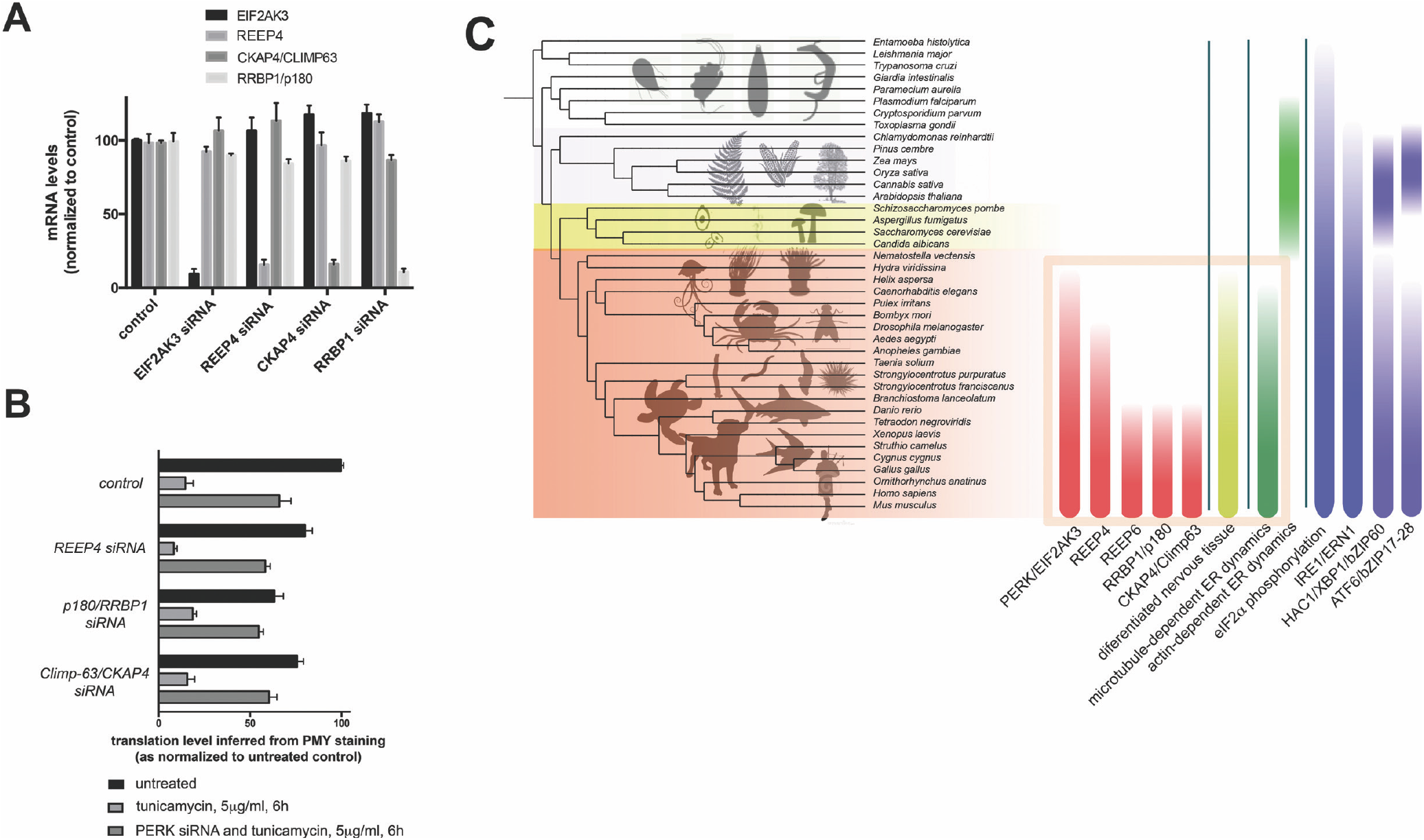
**(A)** qRT-PCR analysis of indicated mRNA transcripts across indicated conditions. Data are derived from 3 biological replicates. **(B)** Image-based puromycylation assay across indicated conditions (4 biological replicates, approx. 2000 cells each). Data are expressed as normalized to control untreated condition, which is set as 100%. **(C)** Visual rendering of estimated evolutionary conservation of indicated genes, as related to specific features of ER-cytoskeleton relationship. Evolutionary conservation is derived from InParanoid database, and layered over an evolutionary tree across indicated eukaryotic phila (OrthoDB, University of Geneva).

**Supplementary Figure 5.**
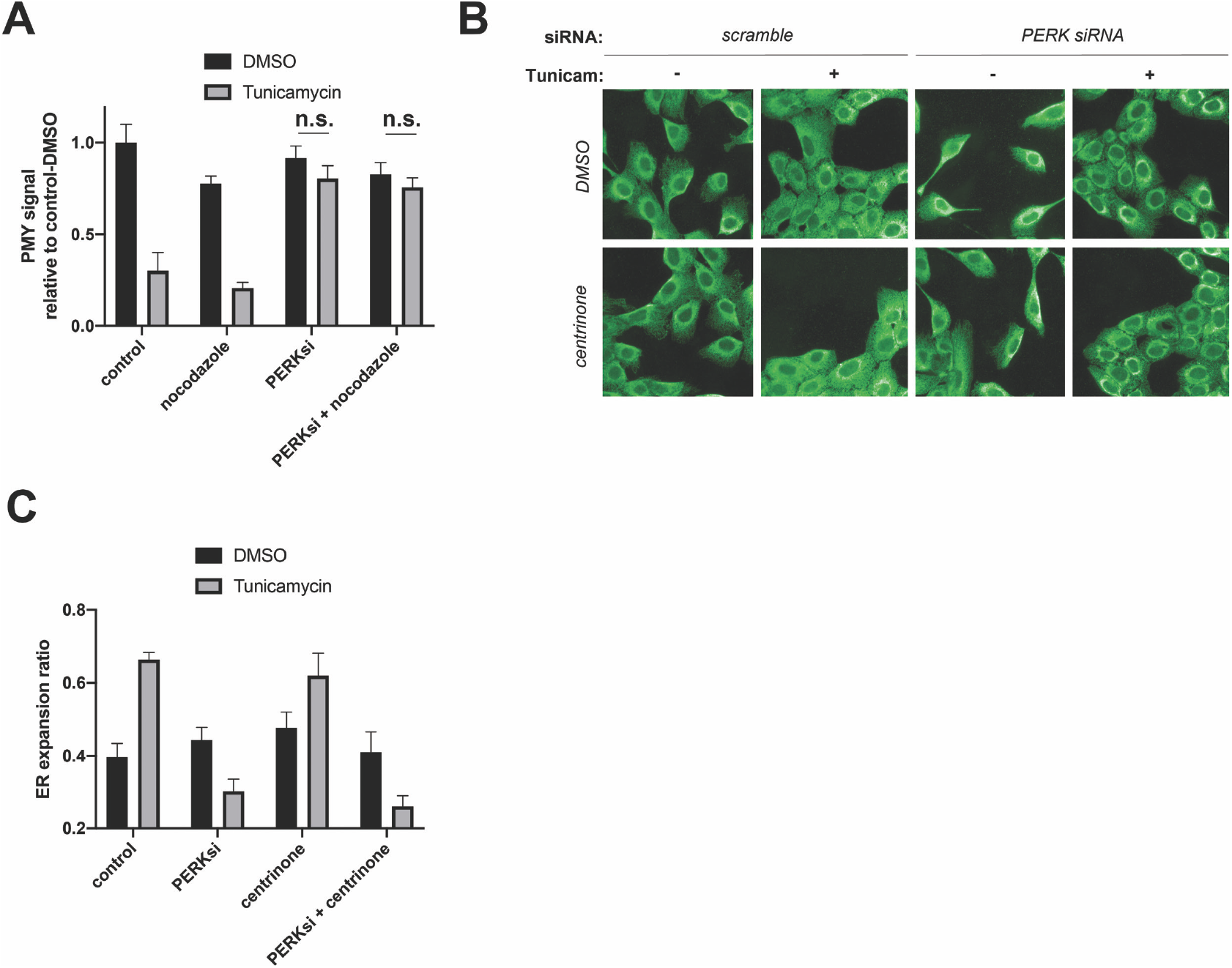
**(A)** Image-based puromycylation assay across indicated conditions (4 biological replicates, approx. 2000 cells each). Data are expressed as normalized to control untreated condition. **(B-C)** ER remodeling assay (immunofluorescence images: calreticulin) across indicated conditions (4 biological replicates, approx. 2000 cells each). Statistical significance across assays was assessed by paired t-Student test; *: *p* <0.05; **: *p* <0.01; ***: *p* <0.005. n.s.: *p* >0.05

**Supplementary Figure 6.**
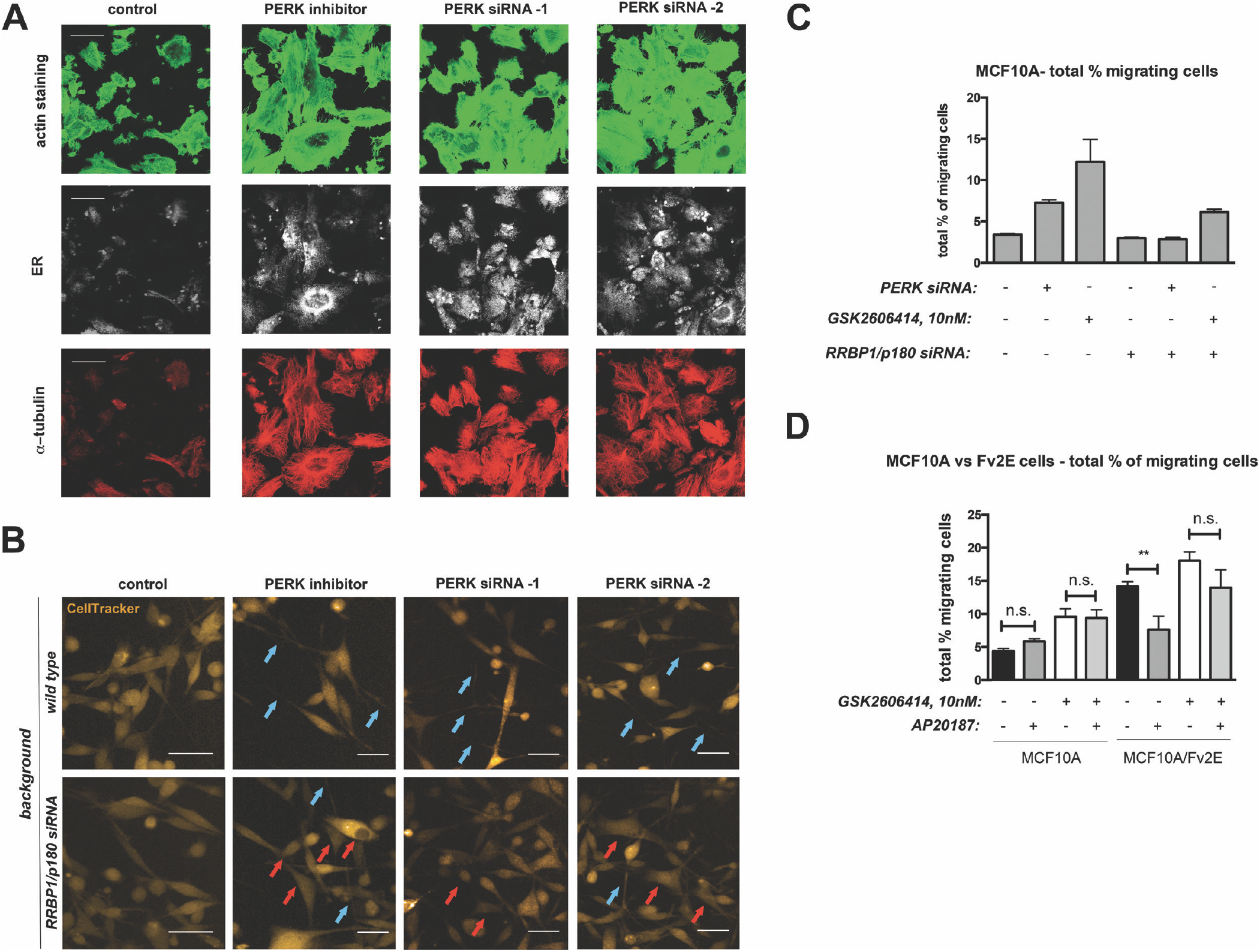
**(A)** Control Transwell™ protrusion formation assay in MCF10A cells across indicated conditions and labels. PERK inhibitor: GSK2606414, 40nM **(B)** Cells subjected to indicated siRNA treatments were plated on soft collagen matrices, counterstained with CellTracker Orange, and imaged by confocal microscopy. Examples of cells with long protrusions (blue arrows), as opposed to cells with no apparent elongated protrusions (red arrows), are indicated. **(C, D)** Cell migration assays across indicated conditions and MCF10A clones, related to experiments shown in Fig. 7F. Data were derived from 6 biological replicates. Statistical significance across assays was assessed by paired t-Student test; *: *p* <0.05; **: *p* <0.01; ***: *p* <0.005. n.s.: *p* >0.05

**Supplementary table 1.**

| PRIMARY ANTIBODIES |  |  |  |
| --- | --- | --- | --- |
| Antigen | Supplier | Reference Num. | Usage |
| Calreticulin | Abcam | ab2907 | Immunofluorescence (1:2000) |
| pSer51 eIF2a | Enzo Biosciences | BML-SA405 | Western Blot (1:1000), IF (1:250) |
| eIF2a | Abgent | P05198 | Western Blot (1:1000) |
| Puromycylated peptide | DSHB | PMY-2A4 | IF (1:2000) |
| HA tag | Roche | 11867423001 | Western Blot (1:1000) |
| α-tubulin | Sigma | T9026 | IF (1:2000) |
| CAMSAP2 | ThermoScientific | PA5-99978 | Western Blot (1:1000) |
| RRBP1 | ThermoScientific | MA5-18302 | IF (1:250) |
| ATF4 | Santa Cruz | sc-390063 | Western Blot (1:1000) |

| siRNA/esiRNA |  |  |
| --- | --- | --- |
| Target | Supplier | Reference |
| PERK | Dharmacon | M-004883-03 |
| PERK | Dharmacon | J-004883-09 |
| PERK | Dharmacon | J-004883-10 |
| PERK | Dharmacon | J-004883-11 |
| PERK | Dharmacon | J-004883-12 |
| PERK | Dharmacon | L-004883-00 |
| PERK | Sigma | EHU030881 |
| IRE1 | Dharmacon | M-004951-02 |
| ATF6 | Dharmacon | M-009917-01 |
| ATL1 | Dharmacon | M-010946-02 |
| RTN4 | Dharmacon | M-010721-00 |
| RTN1 | Dharmacon | M-014138-00 |
| ATL2 | Dharmacon | M-014047-00 |
| ATL3 | Dharmacon | M-010656-00 |
| SREBP1 | Dharmacon | M-006891-01 |
| CKAP4 | Dharmacon | L-012755-01 |
| RRBP1 | Dharmacon | L-011891-02 |
| ATF4 | Dharmacon | L-005125-00 |
| REEP4 | Dharmacon | L-016343-02 |
| REEP5 | Dharmacon | L-019467-01 |
| REEP6 | Dharmacon | L-015555-02 |
| CAMSAP2 | Dharmacon | L-022091-00 |
| CAMSAP2 | Sigma | EHU134911 |

